# Spatial Confidence Sets for Raw Effect Size Images

**DOI:** 10.1101/631473

**Authors:** Alexander Bowring, Fabian Telschow, Armin Schwartzman, Thomas E. Nichols

## Abstract

The mass-univariate approach for functional magnetic resonance imagery (fMRI) analysis remains a widely used and fundamental statistical tool within neuroimaging. However, this method suffers from at least two fundamental limitations: First, with sample sizes growing to 4, 5 or even 6 digits, the entire approach is undermined by the null hypothesis fallacy, i.e. with sufficient sample size, there is high enough statistical power to reject the null hypothesis everywhere, making it difficult if not impossible to localize effects of interest. Second, with any sample size, when cluster-size inference is used a significant *p*-value only indicates that a cluster is larger than chance, and no notion of spatial uncertainty is provided. Therefore, no perception of confidence is available to express the size or location of a cluster that could be expected with repeated sampling from the population.

In this work, we address these issues by extending on a method proposed by *Sommerfeld, Sain, and Schwartzman (2018)* to develop spatial Confidence Sets (CSs) on clusters found in thresholded raw effect size maps. While hypothesis testing indicates where the null, i.e. a raw effect size of zero, can be rejected, the CSs give statements on the locations where raw effect sizes exceed, and fall short of, a *non-zero* threshold, providing both an upper and lower CS.

While the method can be applied to any parameter in a mass-univariate General Linear Model, we motivate the method in the context of BOLD fMRI contrast maps for inference on percentage BOLD change raw effects. We propose several theoretical and practical implementation advancements to the original method in order to deliver an improved performance in small-sample settings. We validate the method with 3D Monte Carlo simulations that resemble fMRI data. Finally, we compute CSs for the Human Connectome Project working memory task contrast images, illustrating the brain regions that show a reliable %BOLD change for a given %BOLD threshold.

## 1. Introduction

Over the last three decades, the Statistical Parametric Mapping procedure (Friston et al., 1994a) for inference of task-fMRI data has prevailed as the international standard within the field of neuroimaging. Incorporating a mass-univariate statistical approach, functional data at each voxel is described in terms of experimental conditions and residual variability included as parameters in a general linear model. From this model, a group-level Statistical Parametric Map (SPM) of *t*-statistic’s contrasting a specified experimental condition relative to a baseline condition is formed. Using a corrected significance level based on the theory of random fields to account for the multiple-comparison problem (Friston et al., 1994b), hypotheses are tested at each voxel independently and the SPM is finally thresholded to localize brain function. While simple by nature, this technique has proven immensely powerful and provided us with the tools to gain deep insight into cognitive function.

There is, however, information that *is not* captured using the current fMRI approach to inference. Specifically, for clusterwise inference, the cluster-level *p*-value only conveys information about a cluster’s spatial extent under the null-hypothesis. Since no information is provided regarding the statistical significance of each voxel comprising a significant cluster, the most we can say is that significant activation has occurred *somewhere* inside the cluster (Woo et al., 2014). An implication of this is that when a large, sprawling cluster covers many anatomical regions, the precise spatial specificity of the activation is in fact poor. While a recent effort has attempted to solve this problem by ‘drilling down’ to find the exact source of activation (Rosenblatt et al., 2018), this can come at the cost of lower statistical power. A related problem of cluster inference is that no information is provided about the spatial variation of significant clusters. For example, if a given fMRI study were to be repeated many times with new sets of subjects, there would of course be variation in the size and shape of clusters found, yet the current statistical results have no way to characterize this variability.

A more pressing issue, perhaps, stems from an age-old paradox caused by the ‘fallacy of the null hypothesis’ (Rozeboom, 1960). The paradox is that while statistical models conventionally assume mean-zero noise, in reality all sources of noise will *never* cancel, and therefore improvements in experimental design will eventually lead to statistically significant results. Thus, the null-hypothesis will, eventually, *always* be rejected (Meehl, 1967). The recent availability of ambitious, large-sample studies (e.g Human Connectome Project (HCP), N=1,200; UK Biobank, N=30,000 and counting) have exemplified this problem. Analysis of high-quality fMRI data acquired under optimal noise conditions has been shown to display almost universal activation across the entire brain after hypothesis testing, even with stringent correction (Gonzalez-Castillo et al., 2012).

For the reasons discussed above, alongside further concerns about misconceptions and the misuse of *p*-values in statistical testing (Nuzzo, 2014, Wasserstein et al., 2016), there has been a growing consensus among sections of the neuroimaging community that the statistical results commonly reported in the literature should be supplemented by effect estimates (Chen et al., 2017, Nichols et al., 2017). The main argument put forward supporting raw effect sizes is that they increase interpretability of the statistical results, highlighting the magnitude of statistically significant differences and providing another layer of evidence to support the overall scientific conclusions inferred from an fMRI study. This may also help tackle reproducibility concerns that have become prominent within the field due to failed attempts in replicating published neuroimaging results based off of statistical testing methods (Poldrack et al., 2017).

In this work, we seek to address all of these issues by applying and extending a spatial inference method initially proposed by *Sommerfeld, Sain, and Schwartzman (2018) (SSS)* to obtain precise confidence statements about where activation occurs in the brain. Unlike hypothesis testing, our spatial Confidence Sets (CSs) allow for inference on *non-zero* raw effect sizes. While the method can be applied to any parameter in a mass-univariate General Linear Model, here we will focus inference on the mean percentage BOLD change raw effect. For a cluster-forming threshold *c*, and a predetermined confidence level 1 − α, the CSs comprise of two sets: the upper CS (denoted 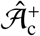, red voxels in Fig. 1), giving all voxels we can assert have a percentage BOLD raw effect size truly *greater* than *c*; and the lower CS (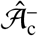, blue voxels overlapped by yellow and red in Fig. 1), for which all voxels *outside* this set we can assert have a percentage BOLD raw effect size truly *less* than *c*. The upper CS is smaller and nested inside the lower CS, and the assertion is made with (1 − α)100% confidence holding simultaneously for both regions. Fig. 1 provides an illustration of the schematic we will use to display the CSs, also showing the point estimate set (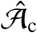, yellow voxels overlapped by red) obtained by thresholding the data at *c*.

**Figure 1:**
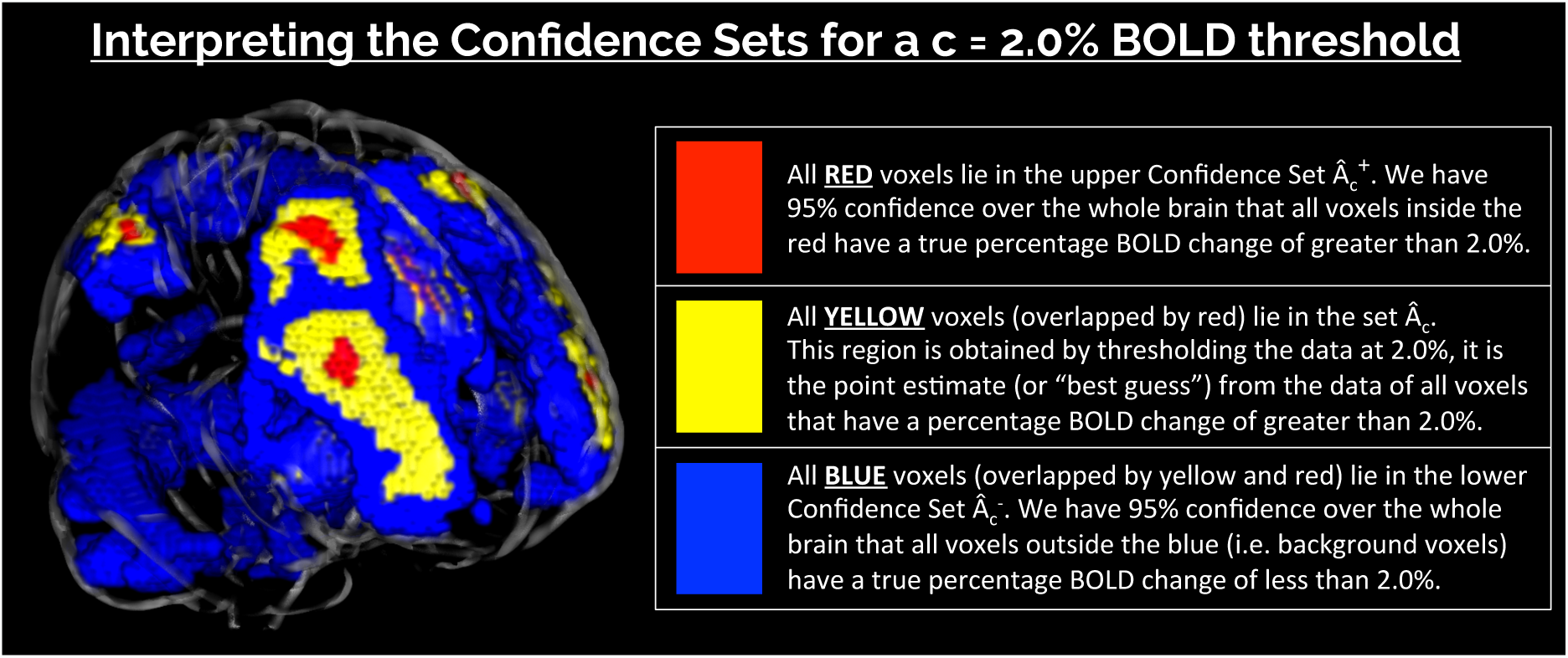
Schematic of the colour-coded regions used to visually represent the Confidence Sets (CSs) and point estimate set. CSs displayed in the glass brain were obtained by applying the method to 80 participants contrast data from the Human Connectome Project working memory task, using a a *c* = 2.0% BOLD change threshold at a confidence level of 1 − α = 95%.

With this interpretation, the CSs can be linked to traditional statistical voxelwise thresholding with control of the familywise error rate (FWE): In a one-sided *t*-test, for the set of level α FWE-significant voxels we have (1 − α)100% confidence that the signal is greater than zero. Put another way, we have (1 − α)100% confidence that the voxelwise level α FWE results are all true positives. The CSs can be viewed as a generalisation of these methods, except that the confidence statement is no longer relative to a signal of zero, but to a non-zero signal magnitude *c*. Users may choose the threshold *c* based on prior knowledge of raw effect sizes reported in previous similar studies to their own; since computation of the CSs is quick, users may also report results for a variety of cluster-forming thresholds as we do in this work.

The motivating data in *SSS* were longitudinal temperature data in North America, and the goal was to infer on areas at risk of climate change. In this work, we are motivated by subject-level fMRI contrast of a parameter estimate maps, and we seek to infer brain areas where a substantial raw effect is present in units of percentage BOLD change. In *SSS*, the CSs were referred to as ‘Coverage Probability Excursion sets’ – shortened to ‘CoPE sets.’

The main contributions of this work are modifications to the *SSS* method for computing CSs that improve the method’s finite-sample performance in the context of neuroimaging. In particular, we propose a combination of the Wild *t*-Bootstrap method and the use of Rademacher variables (instead of Gaussian variables) for multiplication of the bootstrapped residuals, which we find substantially improves performance of the method in moderate sample sizes (e.g. *N* = 60). We also develop a linear interpolation method for computing the boundary over which the bootstrap is applied, and a novel approach for assessing the empirical coverage of the CSs that reduces upward bias in how the simulation results are measured. Another contribution here is that we assess the finite-sample accuracy of the method on synthetic 3D signals that are representative of fMRI activation clusters, whereas *SSS* only considered 2D images. Altogether, we carry out a range of 3D simulations alongside smaller 2D simulations to evaluate our proposed methodological modifications and compare our results to the simulations conducted in *SSS*. Finally, we apply the method to the Human Connectome Project working memory task dataset, operating on the subject-level percentage BOLD change raw effect maps, where we obtain CSs for a variety of cluster-forming thresholds. Here, the method localizes brain activation in cognitive regions commonly associated to working memory, determining with 95% confidence a raw effect of at least 2% BOLD change.

The remainder of this paper is organized as follows. First of all, we summarize the key theory of CSs before detailing our proposed modifications. We then describe the settings used for our simulations, and provide background information about the HCP dataset analyzed in this work. Finally, we report the results of our simulations before presenting the CSs computed for the HCP data.

## 2. Theory

### 2.1. Overview

A comprehensive treatment of the original method, including proofs, can be found in *SSS*. Here we develop the method specifically for the general linear model (GLM) and describe our own enhancements to the method. While the method can be performed for subject-level inference, we will motivate the method in the context of a group-level analysis, describing how the method can be applied to subject-level %BOLD estimate maps in order to obtain group-level CSs making confidence statements about %BOLD effect sizes relating to the entire population from which the participants were drawn.

For a compact domain *S* ⊂ ℝ^*D*^, e.g. *D* = 3, consider the GLM at location ***s*** ∈ *S*,

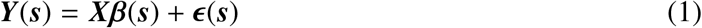

where ***Y***(***s***) is an *N* × 1 vector of observations at ***s***, ***X*** is an *N* × *p* design matrix, β(***s***) is an *p* × 1 vector of unknown coefficients, and **ϵ**(***s***) an *N* × 1 vector of mean-zero errors, independent over observations, and with each ϵ_*i*_ having common variance σ^2^(***s***) and some unspecified spatial correlation. (Throughout we use boldface to indicate a vector-or matrix-valued variable.) In the context of a task-fMRI analysis, ***Y***(***s***) is a vector of subject-level %BOLD response estimate maps obtained by applying a first-level GLM to each of the *N* participants functional data.

For a *p* × 1 contrast vector ***w***, we seek to infer regions of the brain where a contrast of interest ***w**^T^* β has exceeded a fixed threshold *c*. Particularly, we are interested in the noise-free, population cluster defined as:

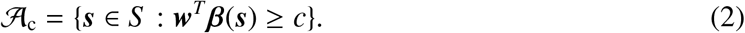

Since we are unable to determine this excursion set in practice, our solution is to find spatial CSs: an upper set 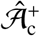 and lower set 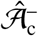 that surround 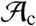 for a desired confidence level of, for example, 95%. We emphasize that these clusters regard the raw units of the signal. Going forward, we assume that the design matrix ***X*** and contrast ***w*** have been carefully chosen so that 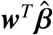 has the interpretation of mean %BOLD change across the group. For example, in a one-sample group fMRI model where data ***Y*** have %BOLD units, choosing ***X*** as a column of 1’s and ***w*** = (1) would ensure that 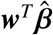 has units of %BOLD change^1^. In this setting, we wish to obtain an upper CS, 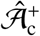, such that we have 95% confidence all voxels *contained* in this set have a population raw effect size *greater* than, for example, c = 2.0% BOLD change, and a lower CS, 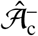, such that we have 95% confidence all voxels *outside* of this set have a population raw effect size *less* than 2.0% BOLD change. Moreover, we desire that the 95% confidence statement holds simultaneously across both CSs at once. *SSS* show that a construction of such CSs is possible within the general linear model framework using the following key result.

**Result 1.***Consider the general linear model setup described in (1). Let* 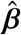 *denote the ordinary least squares estimator of* β, 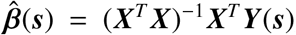, *and define 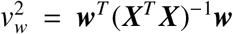 to be the normalised variance of the contrast estimate*.

*Then for a constant k, and for upper and lower CSs defined as*

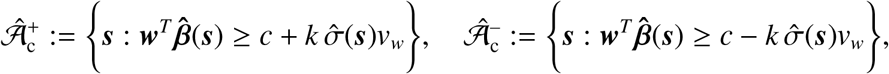

*the limiting coverage of the CSs is*

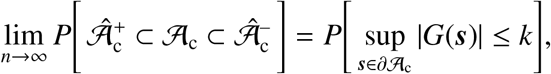

*where* 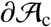 *denotes the boundary of* 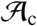*, and G is a smooth Gaussian field on S with mean zero, unit variance, and with the same spatial correlation as each* ϵ_*i*_.

Result 1 is subject to continuity of the relevant fields and some basic conditions on the increments and moments of the error field ϵ. A list of these assumptions, as well as a proof of Result 1, are itemized in *SSS*.

For a pre-determined confidence level 1 − α (e.g. 1 − α = 95%), by choosing *k* such that

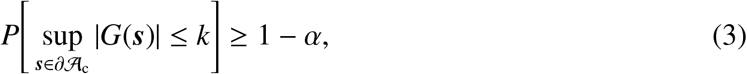

Result 1 ensures with asymptotic probability of 1 − α that 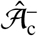 contains the true 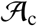, and 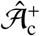 is contained within 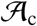. In practice, *k* is determined as the (1 − α)100 percentile of the maximum distribution of the asymptotic absolute error process |*G*(***s***)| over the true boundary set 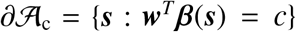. The upper CS taken away from the lower CS 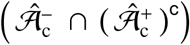 can be interpreted analogously to a standard confidence interval: with a confidence of 1 − α, we can assert the true boundary 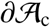 lies within this region. Here, we allude to the classical frequentist interpretation of confidence, where stated precisely, there is a probability of 1 − α that the region 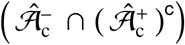 computed from a future experiment fully encompasses the true set boundary 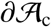.

Application of Result 1 presents us with two challenges: that the boundary set 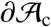 and the critical value *k* are both unknown. To solve the first problem, *SSS* propose using 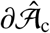 as a plug-in estimate of 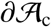. There remain, however, technicalities at to how the boundary is determined in any non-abstract setting, and in particular in a 3D image. In Section 2.3 we propose our own novel method for boundary estimation. Before that, we address the second problem, finding the critical value *k* via a Wild Bootstrap resampling scheme.

### 2.2. The Wild t-Bootstrap Method for Computation of k

To apply Result 1, we require knowledge of the tail distribution of the limiting Gaussian field *G* along the boundary 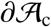. However, the distribution of this field is unknown, because it is dependent on the unknown spatial correlation of the errors ϵ_*i*_. We can approximate the maximum distribution of *G* using the Gaussian Wild Bootstrap (Chernozhukov et al., 2013), also known as the Gaussian Multiplier Bootstrap, which multiplies residuals by random values to create surrogate instances of the random errors.

*SSS* construct *G* as follows: The standardized residuals,

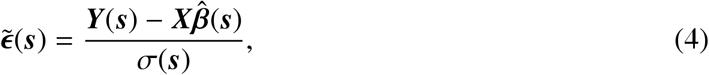

are multiplied by i.i.d. Gaussian random variables, 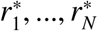, summed and scaled,

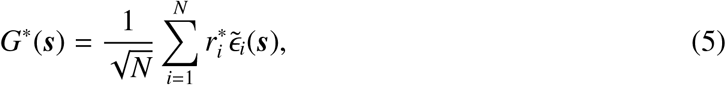

producing a field *G*^∗^ with approximately the same covariance as each error ϵ_*i*_, where the superscript asterisk (∗) indicates these are just one of many bootstrap realizations. With *B* bootstrap samples *G*^∗^, we choose *k* as the (1 − α)100 percentile of the *B* suprema 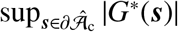 to approximate the LHS of (3) and apply Result 1 to obtain the CSs.

Up to this point, we have summarized the Gaussian Wild Bootstrap methodology as proposed in *SSS*. However, when applying this method to our own simulations, we consistently found that our coverage results fell below the nominal level. This was particularly severe for 3D simulations we conducted using a small sample size (N = 60), where our results in some cases suffered from under-coverage 40% or more below the nominal level (see Fig. 6.2). Hence we made two alterations: First, while *SSS* used Gaussian multipliers, we found improved performance using Rademacher variables, where each *r*_*i*_ takes on 1 or −1 with probability 1/2; others have also reported improved performance with Rademacher variables as well (Davidson and Flachaire, 2008). Second, we implemented a Wild *t*-Bootstrap (Telschow and Schwartzman, 2019) method, normalizing the bootstrapped residuals 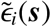 by their standard deviation 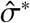. This detail was omitted in the proof of Result 1 provided in *SSS*, where the true standard deviation was assumed to be known. By taking into account the estimation of the standard deviation via the Wild *t*-Bootstrap, we found improved performance for moderate sample sizes. The Wild *t*-Bootstrap version of *G* is

**Figure 2:**
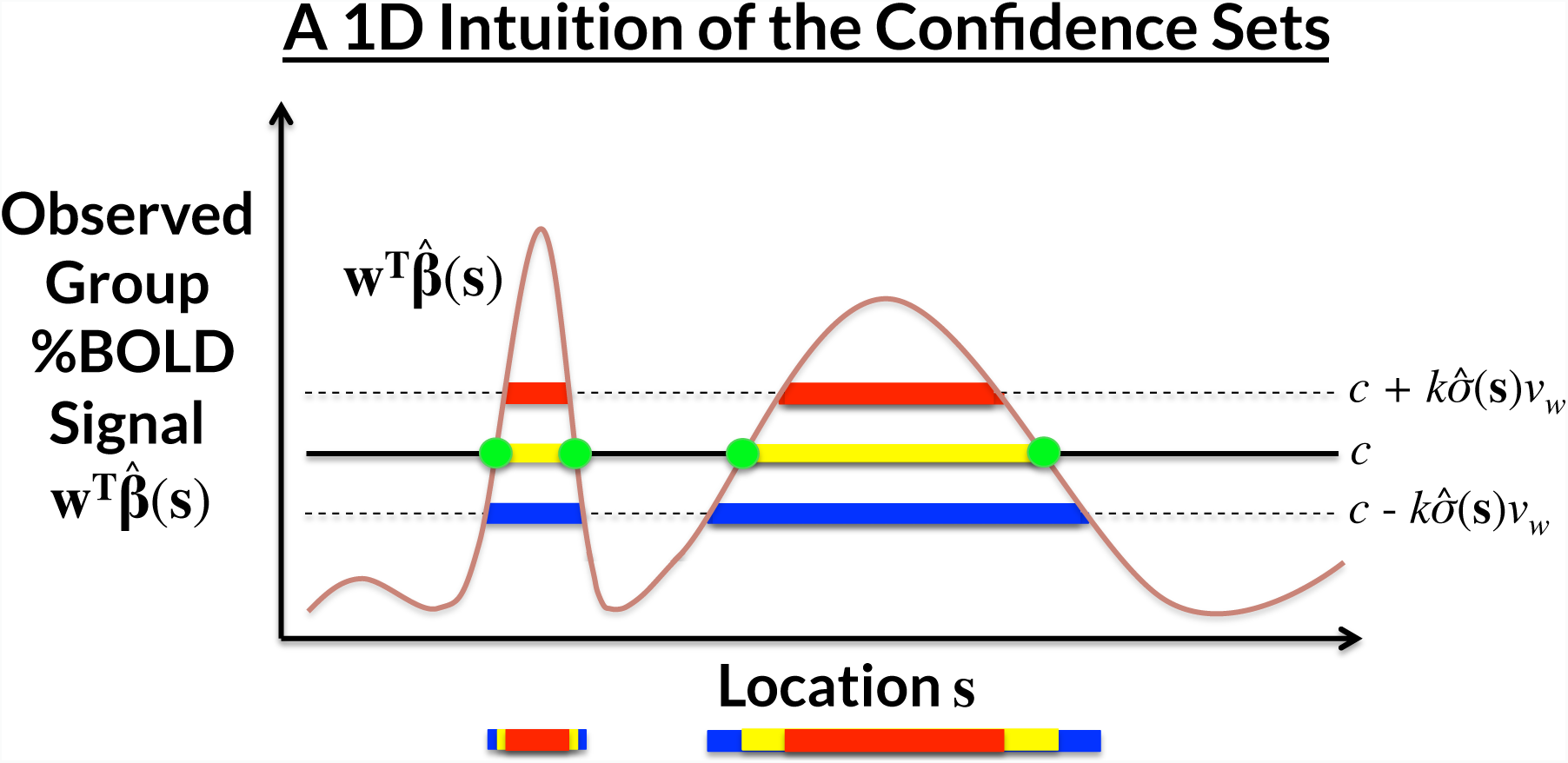
A demonstration of how the CSs are computed for a realization of the GLM ***Y***(***s***) = ***X***β(***s***)+ϵ(***s***) in 1 dimension, for each location ***s***. The yellow voxels 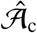 are obtained by thresholding the observed group contrast map at threshold *c*; this is the best guess of 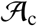, the set of voxels whose true, noise-free raw effect surpasses *c*. The red upper CS 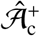 and blue lower CS 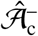 are computed by thresholding the signal at 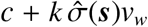 and 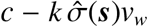, respectively. We have (1 − α)100% confidence that 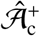 ⊂ 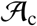 ⊂ 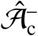, i.e. that 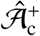 (red) is completely within the true 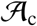, and 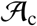 is completely within 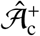 (blue). We find the critical value *k* from the (1 − α)100 percentile of the maximum distribution of the absolute error process over the estimated boundary 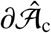(green ●’s) using the Wild *t*-Bootstrap; 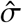 is the estimated standard deviation and *v*_*w*_ is the normalised contrast variance.

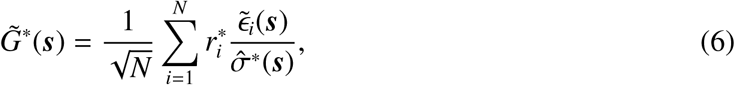

where 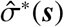 is the standard deviation of the present realization of the bootstrapped residuals 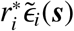. We then determine *k* as described above but using 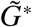 instead of *G*^∗^. Going forward, we refer to this method as the “Wild *t*-Bootstrap”, to be contrasted with the original “Gaussian Wild Bootstrap” method proposed in *SSS*.

With these two alterations we found a dramatic increase in performance for small sample sizes in 3D simulations. Notably, in contrast to the Gaussian Wild Bootstrap, our simulation results presented in Section 4 suggest that empirical coverage rates for this modified procedure remain valid, i.e. stay *above* the nominal level.

### 2.3. Approximating the Boundary on a Discrete Lattice

In the previous section, we described the ideal bootstrap procedure used to obtain the maximum distribution of *G* along the boundary 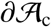 in order to apply Result 1. However, in any practical application, data will be observed on a discrete grid of lattice points at a fixed resolution. Therefore, a key challenge is how to appropriately approximate the true continuous boundary 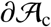 from the lattice representation of the data.

In *SSS*, spline-interpolation was used to estimate a 1D boundary at a resolution greater than their 2D sampled field (*SSS*, Section 4.1). However, to apply the method to fMRI data we will work with 3D images, and estimating a 2D spline boundary for a 3D field is more involved, requiring careful tuning of the spline basis to accommodate the structure of the 3D signal. Instead, we choose to use a first-order weighted linear interpolation method to approximate the signal values at estimated locations along the true, continuous boundary 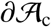, providing a method of boundary estimation that is less computationally intensive than spline interpolation.

Consider two adjacent points on the lattice, ***s_O_*** and ***s_I_***, such that ***s_O_*** lies outside of 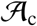, while ***s_I_*** lies inside 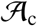. By the definition of 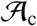, ***w**^T^* β(***s_O_***) < *c*, and ***w**^T^* β(***s_I_***) ≥ *c*. Under the assumption that the component of the signal between ***s_O_*** and ***s_I_*** increases linearly, we can find the location ***s***^∗^ between ***s_O_*** and ***s_I_*** such that ***w**^T^* β(***s***^∗^) = *c*, our estimate of where the true continuous boundary 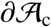 crosses between ***s_O_*** and ***s_I_***. We can then construct a linear interpolant for the location ***s***^∗^, using weights

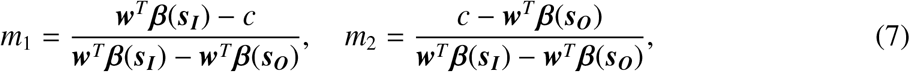

for locations ***s_O_*** and ***s_I_***, respectively. By construction, applying *m*_1_ and *m*_2_ to the contrast image returns the threshold: *m*_1_***w**^T^* β(***s_O_***) + *m*_2_***w**^T^* β(***s_I_***) = ***w**^T^* β(***s***^∗^) = *c*. Applied to standardized residuals 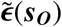 and 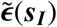, we can likewise obtain the residuals at the estimated continuous boundary point 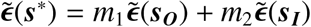.

By repeating this procedure for all adjacent points ***s_O_*** and ***s_I_*** that lie on the lattice either side of 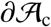, we are able to estimate the standardized residual values at locations that should approximately sample the true continuous boundary 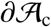, and thus we can apply the ideal bootstrap procedure outlined in Section 2.2. Of course, in practice we apply this interpolation method on the observed, noisy data, using the plug-in estimated boundary 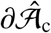.

In the simulation results in Section 4, we assess performance of the method when the bootstrap procedure is carried out over the true boundary 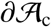, and the plug-in estimated boundary 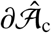 that must be used in practice.

### 2.4. Assessment of Continuous Coverage on a Discrete Lattice

In testing the finite-sample validity of our method through simulation, it is imperative that we are able to accurately measure when violations of the subset condition 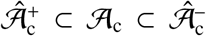 occur. While this may seem a trivial task, as touched on in the previous section, the boundaries of each of these three sets can become ambiguous when data are collected on a discrete lattice.

To illustrate this point, consider the configuration of sets displayed in Fig. 3a. In this instance, suppose the right half of the image corresponds to 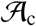 (green pixels overlapped by yellow), and yellow pixels belong to 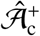. We wish to determine whether the condition 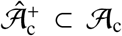 has been violated or not. One may argue that at the resolution for which the data have been acquired, all pixels that belong to 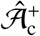 also belong to 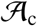, and therefore no violation has occurred. However, the example presented in Fig. 3a has in fact been derived from a 2D simulation conducted at a higher resolution: this 50 × 50 simulation was obtained by down-sampling a 100 × 100 grid by dropping every other pixel. Fig. 3a displays the sets 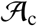 and 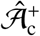 from the down-sampled, low resolution simulation, while Fig. 3b shows the same set of results at the original resolution. In Fig. 3b we see that there *has* been an upcrossing of the yellow pixels belonging to 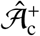 over the boundary of the green, and therefore the subset condition 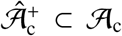 *has* been violated. From this simulation, it is clear that when we conclude that no violation has occurred in situations like Fig. 3a, our empirical coverage will miss violations and be positively biased. By an analogous argument the same issue occurs when testing violations of 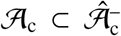.

**Figure 3:**
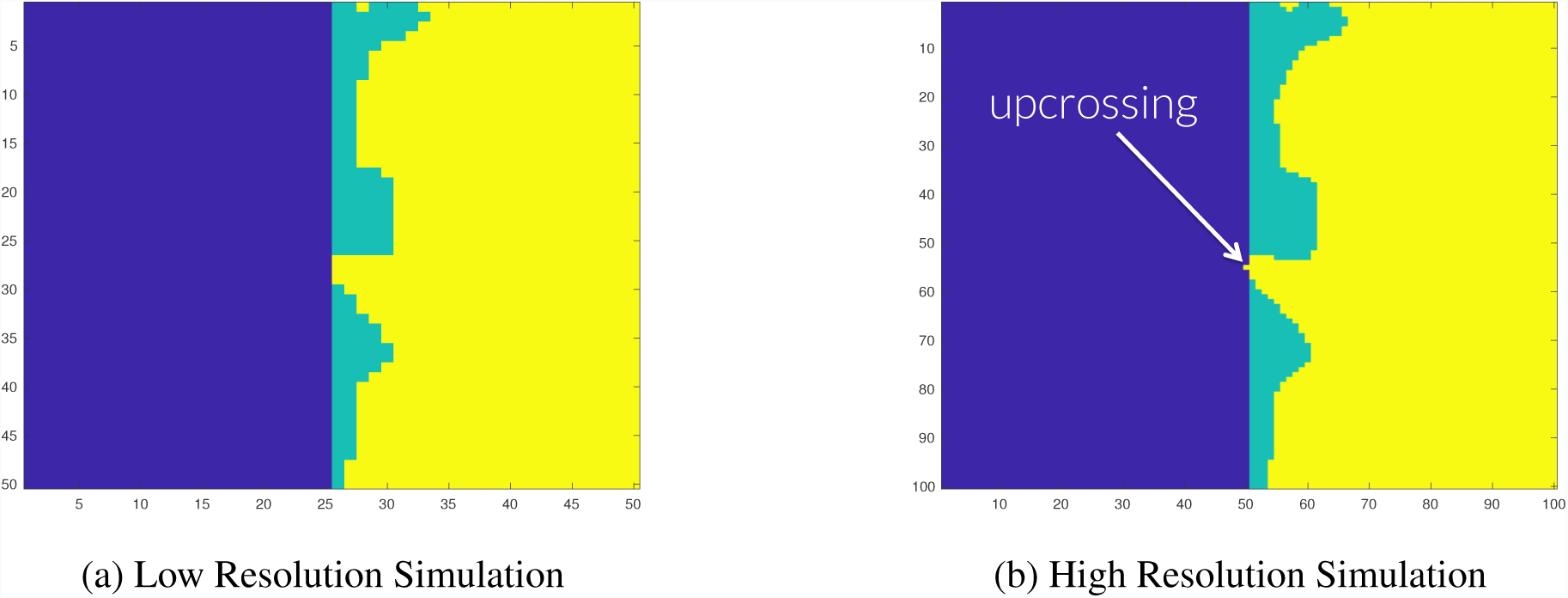
Demonstrating the resolution issue for testing the subset condition 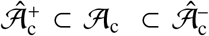 **Figure 3a**: Here 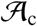 is comprised of the right half of the image (all green and yellow pixels), and 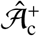 is shown as yellow pixels. It appears that 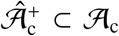. **Figure 3b**: The same configuration as Fig. 3a at double the resolution. Here, we have enough detail to see that 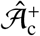 has crossed the boundary 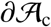 (yellow seeping into blue), and the subset condition 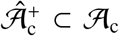 has been violated.

In *SSS* this direct comparison of the lattice representation of the three sets was used to assess coverage in the simulations. While they observed this phenomenon of missed violations leading to over-coverage, the proposed solution was to sequentially increase the resolution of the data. We instead again make use of interpolation.

Since, in simulation, we know the true continuous mean image and 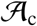, following the method described in Section 2.2 we can obtain weights *m*_1_ and *m*_2_ to interpolate between points ***s_O_*** and ***s_I_*** either side of the true, continuous boundary 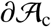, in order to find a location ***s***^∗^ that approximately lies on the boundary (if the true mean is linear, it would be exactly on the boundary). To determine if 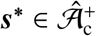, we then re-apply the weights *m*_1_ and *m*_2_ and assess whether

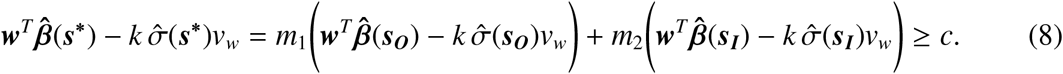

If the inequality holds, then by definition 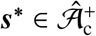. Otherwise, 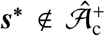, and therefore we can conclude that the subset condition 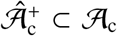 has been violated. By checking whether 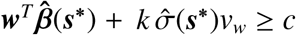, we can similarly test for a violation of 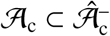.

By applying this interpolation scheme to all pairs of lattice points with one point inside, one outside, the lattice representation of the boundary, we have devised a method to more accurately assess violations of the subset condition 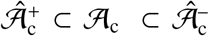 for configurations similar to Fig. 3a. We applied this method for testing the subset condition in our simulations alongside a direct comparison of the lattice representations of the three sets of interest as was done in *SSS*. The addition of the weighted interpolation method caused a considerable decrease in the empirical coverage results towards the nominal level in all of our 3D simulations. Using the direct comparison of the three sets on its own here essentially determined total empirical coverage (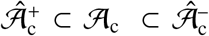 for all simulation runs), even when using small sample sizes and a low nominal coverage level. This is likely to be because the discrete lattice of observed data points is relatively less dense inside the true continuous process for larger, 3D settings, and therefore more violations of the subset condition are missed if only a direct comparison of the lattice representation of the CSs is carried out.

## 3. Methods

### 3.1. Simulations

In this section we describe the settings used in order to evaluate the CSs obtained for synthetic data. As a simplified instance of the general linear model setup described in Section 2.1, we simulate 3000 independent samples of the signal-plus-noise model

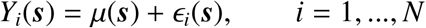

using a range of signals µ(***s***), Gaussian noise structures ϵ_*i*_(***s***) with stationary and non-stationary variance, in two- and three-dimensional regions *S*. We compute the critical value *k*, applying the Wild *t*-Bootstrap method outlined in Section 2.2 with *B* = 5000 bootstrap samples to both the true boundary 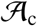 and the plug-in boundary 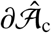 that would be used in practice. The boundaries were obtained using the interpolation method outlined in Section 2.3. We then compare the empirical coverage – the percentage of trials that the true thresholded signal is completely contained between the upper and lowers CSs (i.e. the number of times for which 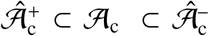) – across the two sets of results, using the assessment method outlined in Section 2.4. In each simulation, we apply the method for sample sizes of N = 60, 120, 240 and 480, and using three nominal coverage probability levels 1 − α = 0.80, 0.90 and 0.95.

### 3.2. 2D Simulations

We analyzed the performance of the CSs on a square region of size 100 × 100. For the true underlying signal µ(***s***) we considered two different raw effects: First, a linear ramp that increased from a magnitude of 1 to 3 in the x-direction while remaining constant in the y-direction (Fig. 4.1a). Second, a circular effect, created by placing a circular phantom of magnitude 3 and radius 30 in the centre of the search region, which was then smoothed using a 3 voxel FWHM Gaussian kernel (Fig. 4.1b). If we were to assume that each voxel had a size of 2mm^3^, we note that this would amount to applying smoothing with a 6mm FWHM kernel, a fairly typical setting used in fMRI analyses.

**Figure 4.1:**
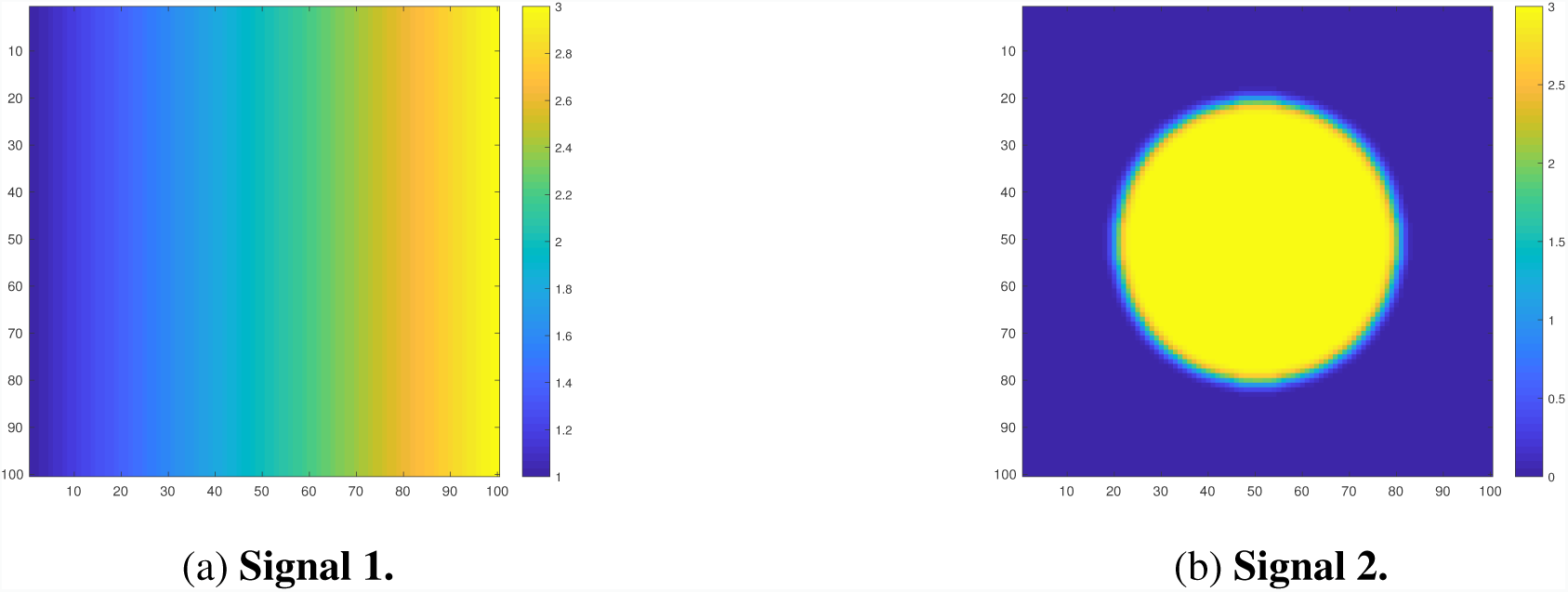
Linear ramp and circular signals µ(***s***) **Figure 4.1a: Signal 1.** A linear ramp signal that increases from magnitude of 1 to 3 in the x-direction. **Figure 4.1b: Signal 2.** A circular signal with magnitude of 3 and radius of 30, centred within the region and convolved with a 3 voxel FWHM Gaussian kernel.

**Figure 4.2:**
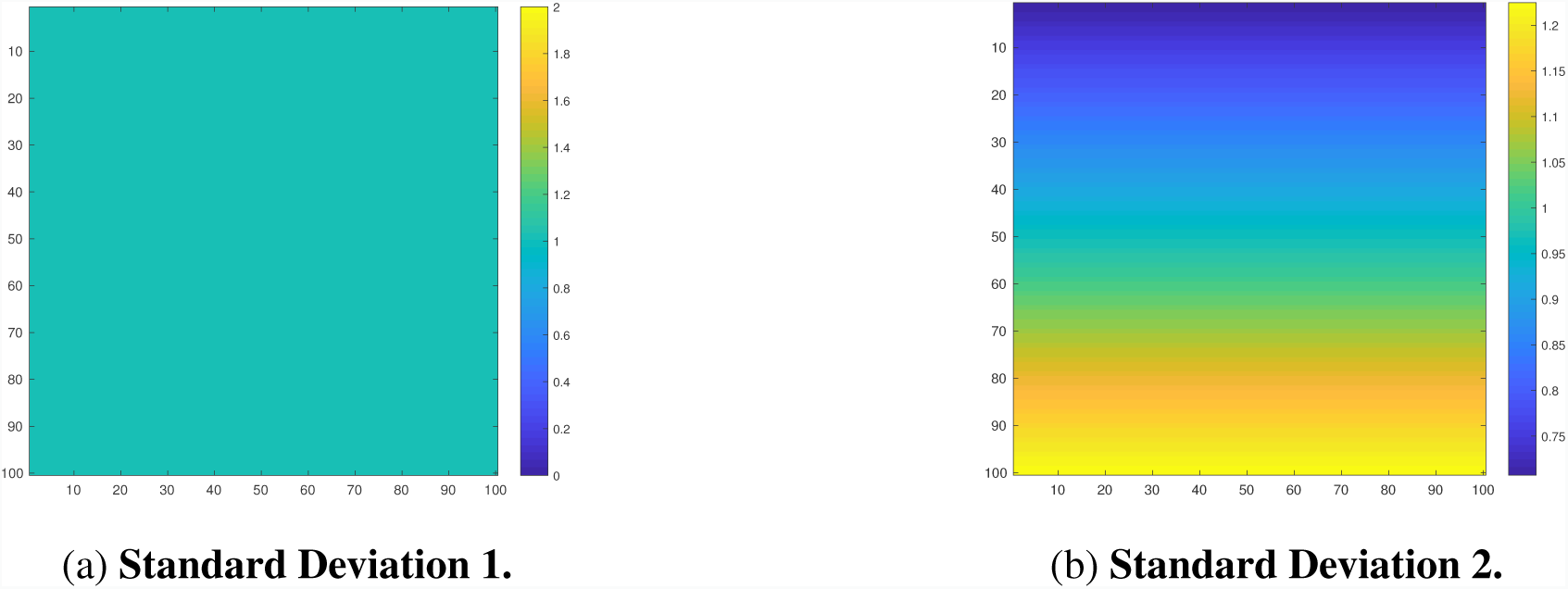
Stationary and non-stationary standard deviation fields of the noise E_*i*_(***s***). **Figure 4.2a: Standard Deviation 1.** Stationary variance of 1 across the region. **Figure 4.2b: Standard Deviation 2.** Non-stationary (linear ramp) standard deviation field increasing from 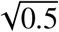 to 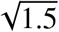 in the y-direction.

To each of these signals we added subject-specific Gaussian noise ϵ_*i*_, also smoothed using a 3 voxel FWHM Gaussian kernel, with homogeneous and non-homogeneous variance structures:

The first noise field had a spatially constant standard deviation of 1 (Fig. 4.2a), the second field had a linearly increasing standard deviation structure in the y-direction from 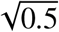 to 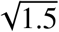 while remaining constant in the x-direction (Fig. 4.2b). Thus, the variance of this noise field spatially increased in the y-direction from 0.5 to 1.5 in a non-linear fashion.

Altogether, the two underlying signals and two noise sources gave us four separate trials; across all of the simulations, we obtained Confidence Sets for the noise-free cluster 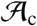 at a cluster-forming threshold of *c* = 2.

### 3.3. 3D Simulations

Four signal types µ(***s***) were considered to analyze performance of the method in three dimensions. The first three of these signals were generated synthetically on a cubic region of size 100 × 100 × 100: Firstly, a small spherical effect, created by placing a spherical phantom of magnitude 3 and radius 5 in the centre of the search region, which was then smoothed using a 3 voxel FWHM Gaussian kernel (Fig. 5a). Secondly, a larger spherical effect, generated identically to the first effect with the exception that the spherical phantom had a radius of 30 (Fig. 5b). Lastly, we created an effect by placing four spherical phantoms of magnitude 3 in the region of varying radii and then smoothing the entire image using a 3 voxel FWHM Gaussian (Fig. 5c). For each of these signals, the final image was re-scaled to have a maximum intensity of 3.

**Figure 5:**
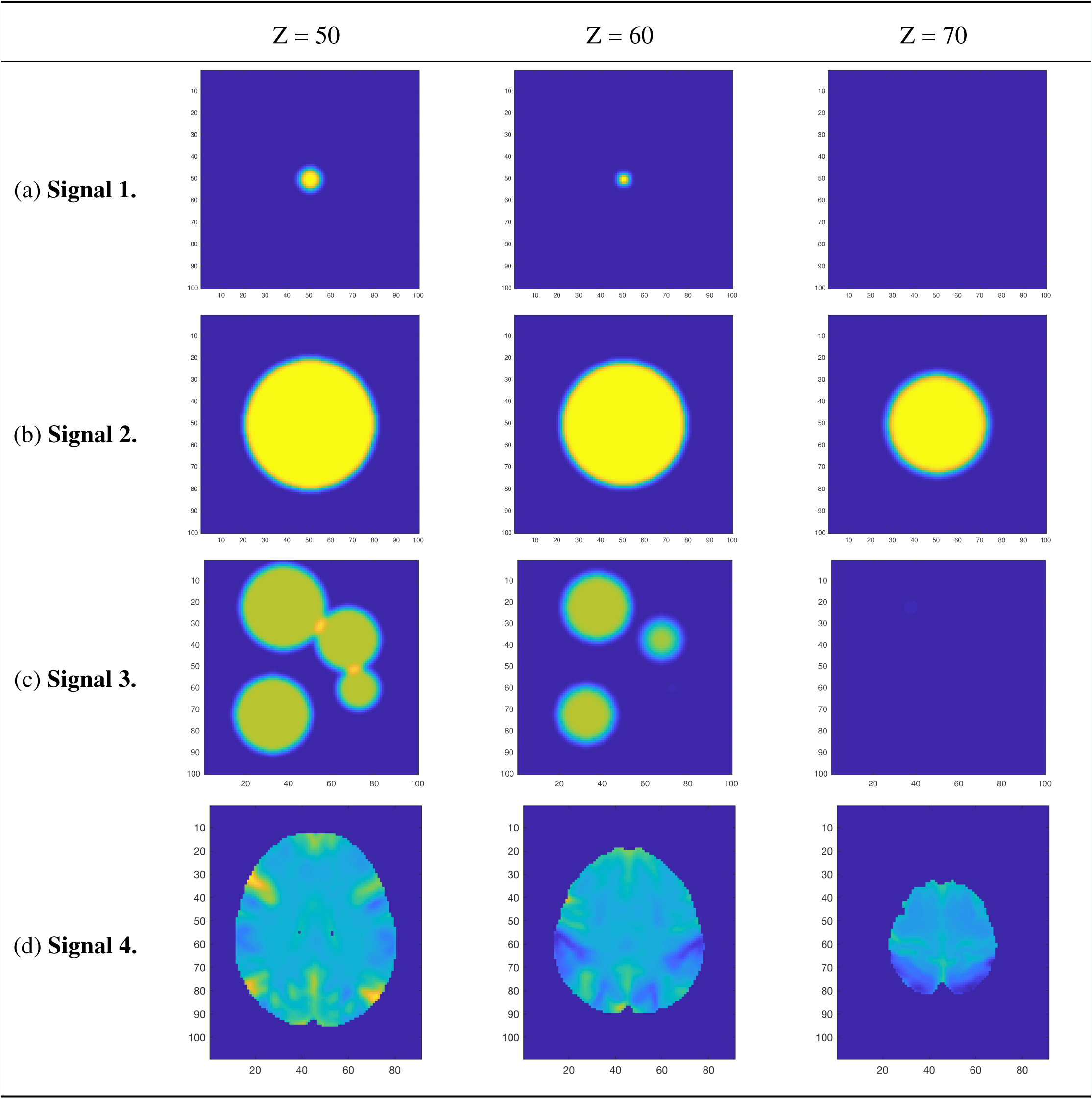
The four 3D signal types µ(***s***), from top-to-bottom: small sphere, large sphere, multiple spheres, and the UK Biobank full mean image. Note that the colormap limits for the first three signal types are from 0 to 3, while the colormap limits for the UK Biobank mean image is from −0.4 to 0.5.

Similar to the two-dimensional simulations, for the three signals described above we added 3-voxel smoothed Gaussian noise of homogeneous and heterogeneous variance structures. The first noise field had a spatially constant standard deviation of 1, while the second field had a linearly increasing standard deviation in the z-direction from 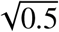 to 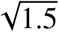, while remaining constant in both the x- and y-directions. For all three effects, we obtained Confidence Sets for the threshold *c* = 2.

For the final signal type, we took advantage of big data that has been made available through the UK Biobank in an attempt to generate an effect that replicated the true %BOLD change induced during an fMRI task. We randomly selected 4000 subject-level contrast of parameter estimate result maps from the Hariri Faces/Shapes task-fMRI data collected as part of the UK Biobank brain imaging study. Full details on how the data were acquired and processed is given in *Miller et al. (2016)*, *Alfaro-Almagro et al. (2018)* and the UK Biobank Showcase; information on the task paradigm is given in *Hariri et al. (2002)*. From these contrast maps, we computed a group-level full mean (Fig. 5d) and full standard deviation image. In the final simulation, we used the group-level Biobank mean image as the true underlying signal µ(***s***) for each subject, and the full standard deviation image was used for the standard deviation of each simulated subject-specific Gaussian noise field ϵ_*i*_(***s***) added to the true signal. Because of the considerably large sample size of high-quality data from which these maps have been obtained, we anticipate that both of these images are highly representative of the true underlying fields that they approximate. Both images were masked using an intersection of all 4000 of the subject-level brain masks.

Once again, we smoothed the noise field using a 3 voxel FWHM Gaussian kernel; since the Biobank maps were written with voxel sizes of 2mm^3^, this is analogous to applying 6mm FWHM smoothing to the noise field of the original data. We obtained Confidence Sets for a threshold of *c* = 0.25% BOLD change.

### 3.4. Application to Human Connectome Project Data

For a real-data demonstration of the method proposed here, we computed CSs on 80 participants data from the Unrelated 80 package released as part of the Human Connectome Project (HCP, S1200 Release). We applied the method to subject-level contrast maps obtained for the 2-back vs 0-back contrast from the working memory task results included with the dataset. To compare the CSs with results obtained from standard fMRI inference procedures, we also performed a traditional statistical group-level analysis on the data. A one-sample *t*-test was carried out in SPM, using a voxelwise FWE-corrected threshold of *p* < 0.05 obtained via permutation test with SPM’s SnPM toolbox. We chose to use the HCP for its high-quality task-fMRI data, the working memory task specifically picked for its association with cognitive activations in sub-cortical networks that can not be distinguished by the anatomy. Full details of the task paradigm, scanning protocol and analysis pipeline are given in *Barch et al. (2013)* and *Glasser et al. (2013)*, here we provide a brief overview.

For the working memory task participants were presented with pictures of places, tools, faces and body parts in a block design. The task consisted of two runs, where on each run a separate block was designated for each of the image categories, making four blocks in total. Within each run, for half of the blocks participants undertook a 2-back memory task, while for the other half a 0-back memory task was used. Eight EVs were included in the GLM for each combination of picture category and memory task (e.g. 2-back Place); we compute CSs on the subject-level contrast images for the 2-back vs 0-back contrast results that contrasted the four 2-back related EVs to the four 0-back EVs.

Imaging was conducted on a 3T Siemans Skyra scanner using a gradient-echo EPI sequence; TR = 720ms, TE = 33.1 ms, 208 × 180 mm FOV, 2.0 mm slice thickness, 72 slices, 2.0 mm isotropic voxels, and a multi-band acceleration factor of 8. Preprocessing of the subject-level data was carried out using tools from FSL and Freesurfer following the ‘fMRIVolume’ HCP Pipeline fully described in *Glasser et al. (2013)*. To summarize, the fundamental steps carried out to each individual’s functional 4D time-series data were gradient unwarping, motion correction, EPI distortion correction, registration of the functional data to the anatomy, non-linear registration to MNI space (using FSL’s Non-linear Image Registration Tool, FNIRT), and global intensity normalization. Spatial smoothing was applied using a Gaussian kernel with a 4mm FWHM.

Modelling of the subject-level data was conducted with FSL’s FMRIB’s Improved Linear Model (FILM). The eight working task EVs were included in the GLM, with temporal derivatives terms added as confounds of no interest, and regressors were convolved using FSL’s default double-gamma hemodynamic response function. The functional data and GLM were temporally filtered with a high pass frequency cutoff point of 200s, and time series were prewhitened to remove autocorrelations from the data.

In comparison to a typical fMRI study, the 4mm FWHM smoothing kernel size used in the HCP preprocessing pipeline is modest. Because of this, we applied additional smoothing to the final contrast images to emulate maps smoothed using a 6mm FWHM Gaussian kernel.

## 4. Results

### 4.1. Methodological Comparisons

In this work we have proposed two fundamental methodological changes to the procedures carried out in *SSS*: in Section 2.2 we suggested the Wild *t*-Bootstrap instead of the Gaussian Wild Bootstrap used for *SSS*, and in Section 2.4 we introduced the interpolation method for assessing empirical coverage alongside the direct comparison methods used for *SSS*. Here, we show the impact of these methodological innovations on the empirical coverage results from simulations carried out using two different synthetic signals, the 2D circular signal (**Signal 2.** in Fig. 4.1b) and the 3D large spherical signal (**Signal 2.** in Fig. 5). The standard deviation of the subject-specific Gaussian noise fields ϵ_*i*_(***s***) had a stationary variance of 1 across the region in both simulations (for the 2D case, this corresponds to **Standard Deviation 1.** in Fig. 4.2).

Empirical coverage results for each of the three confidence levels 1 − α = 0.80, 0.90 and 0.95 are presented for the 2D circular signal in Fig. 6.1 and for the 3D large spherical signal in Fig. 6.2. In both simulations, for all methods the bootstrap procedure was carried out over the estimated boundary 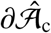 (as must be done with real data). In each figure, the green curves highlight the results for the Gaussian Wild Bootstrap and coverage assessment method that were applied in *SSS*. The red curves highlight the results for the Wild *t*-Bootstrap and interpolation assessment method that we have proposed.

**Figure 6.1:**
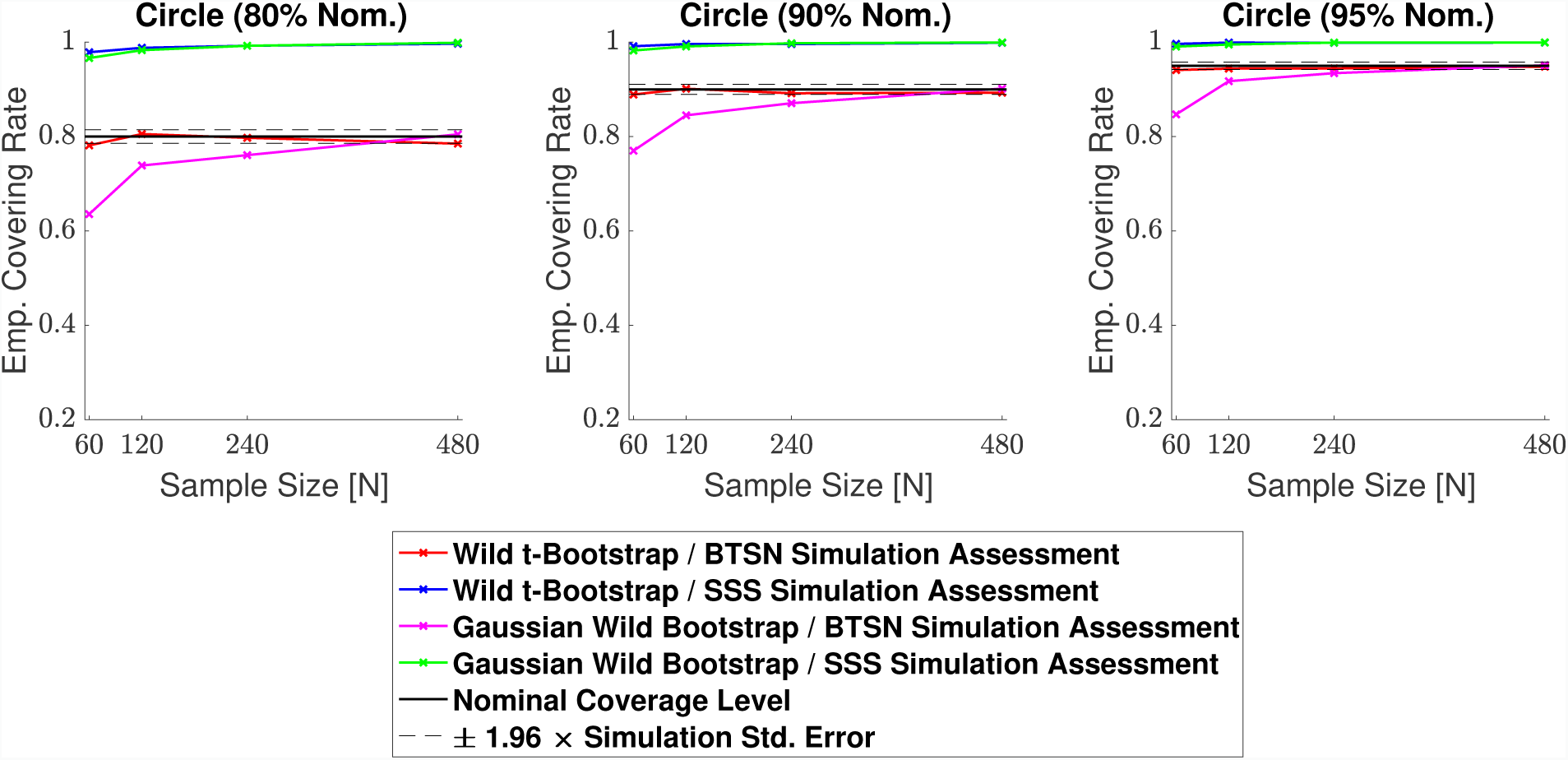
Coverage results for the 2D circular signal simulation with homogeneous Gaussian noise (**Signal 2.**, **Standard deviation 1**. in Fig. 4.2). Empirical coverage results are presented for implementations of the CS method with and without the Wild t-Bootstrap we propose in Section 2.2 and the interpolation schema for assessing simulations results we propose in Section 2.4. All empirical coverage results for simulations using the *SSS* assessment method are close to 100%, suggesting that this assessment substantially biases the results upwards. Using our proposed assessment method, while both the Wild *t*-Bootstrap and Gaussian Wild bootstrap converge to the nominal level, the Wild *t*-Bootstrap performed better for small sample sizes.

**Figure 6.2:**
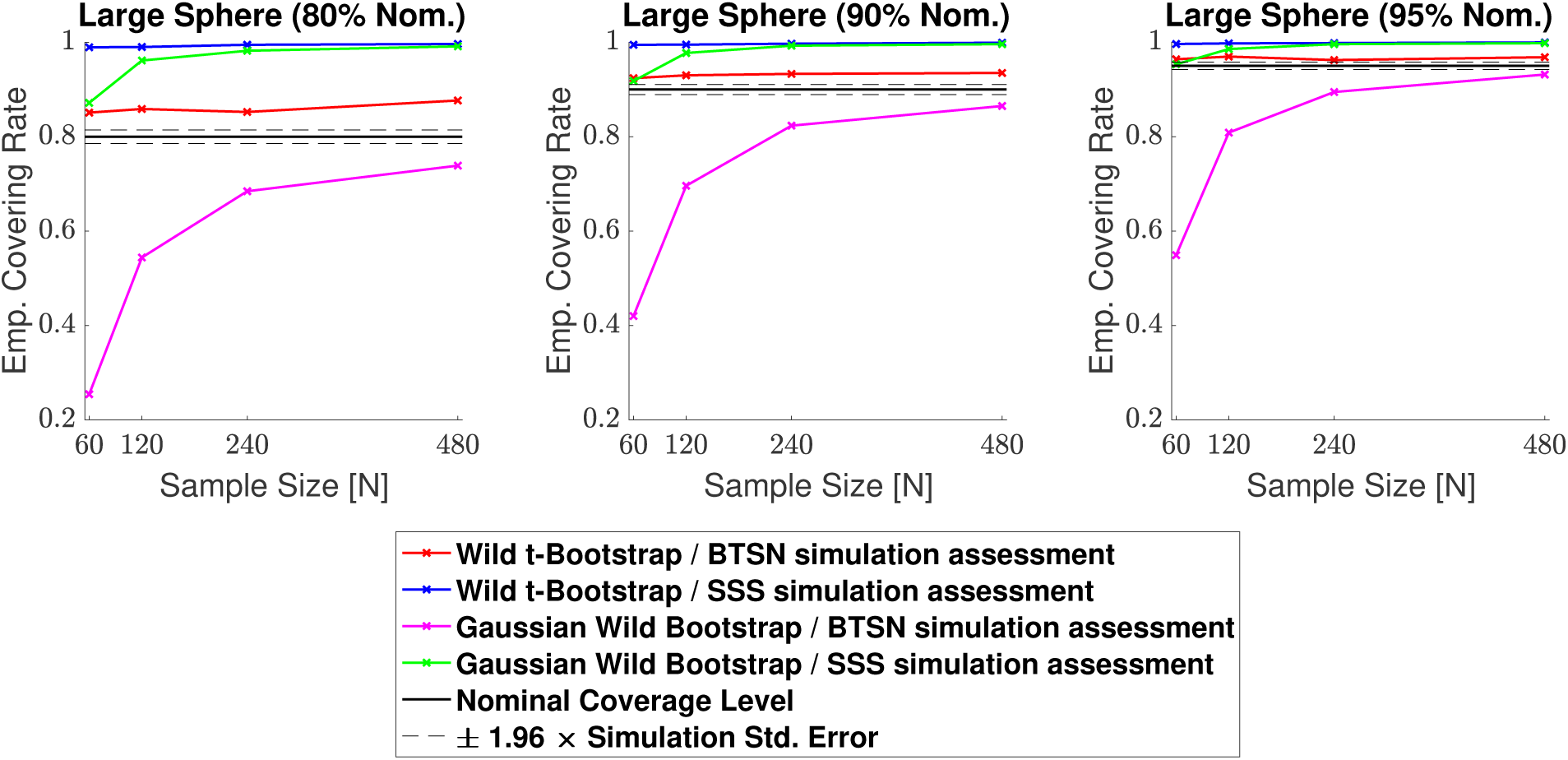
Coverage results for the 3D large spherical signal (**Signal 2**. in Fig. 5) simulation with homogeneous Gaussian noise. Empirical coverage results are presented for implementations of the CS method with and without the Wild *t*-Bootstrap we propose in Section 2.2, and the interpolation schema for assessing simulations results we propose in Section 2.4. Once again, all simulations using the *SSS* assessment method quickly converge to close to 100%. Using our proposed assessment method, the Gaussian Wild bootstrap had severe under-coverage for small sample sizes, while the Wild *t*-Bootstrap results hover slightly above the nominal level for all sample sizes.

In Fig. 6.1 and Fig. 6.2, all simulations using the direct comparison assessment (SSS Simulation Assessment) produced results substantially above the nominal level, converging to almost 100% for both the Gaussian Wild Bootstrap (green curves) and Wild *t*-Bootstrap (blue curves) methods across all three confidence levels. We suspect this is due to the resolution issue described in Section 2.4, suggesting that this assessment method missed violations of the coverage condition 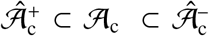 causing a considerable positive bias in all of these results. Further evidence of this is suggested by the empirical coverage obtained for simulations using the interpolation assessment method (BTSN Simulation Assessment, pink and red curves), which appear to be converging much closer to the nominal level as is theoretically expected by Result 1.

Considering only the results using the interpolation assessment, in both figures empirical coverage for the Wild Bootstrap method (pink curves) came below the nominal level for small sample sizes. For the 2D circle simulation, the empirical coverage result for 60 subjects was 84.7% for the nominal target of 1 − α = 0.95 (right plot in Fig. 6.1). For the 3D spherical simulation this under-coverage was even more severe, where the corresponding empirical coverage result was 54.9% (right plot in Fig. 6.2). In comparison, coverage performance for the Wild *t*-Bootstrap method (red curves) was much improved, staying close to the nominal level in both the 2D and 3D simulations across all sample sizes. While for the 3D spherical signal the empirical coverage remained slightly above the nominal target, for the circular signal almost all results lie within the 95% confidence interval of the nominal coverage level. For these reasons, in the remaining simulation results presented in this section we only consider the Wild *t*-Bootstrap method with our proposed interpolation assessment.

### 4.2. 2D Simulations

Empirical coverage results for each of the three confidence levels 1 − α = 0.80, 0.90 and 0.95, are presented for the linear ramp signal (**Signal 1.** in Fig. 4.1a) in Fig. 7.1, and for the circular signal (**Signal 2.** in Fig. 4.1b) in Fig. 7.2. Results are also presented in tabular format in Table. S1. In both plots, results obtained for simulations applying the bootstrap procedure over the estimated boundary 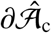 are displayed with a solid line, while results for simulations using the true boundary 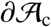 are displayed with a dashed line. We emphasize that when computing CSs for real data, only the estimated boundary can be used.

**Figure 7.1:**
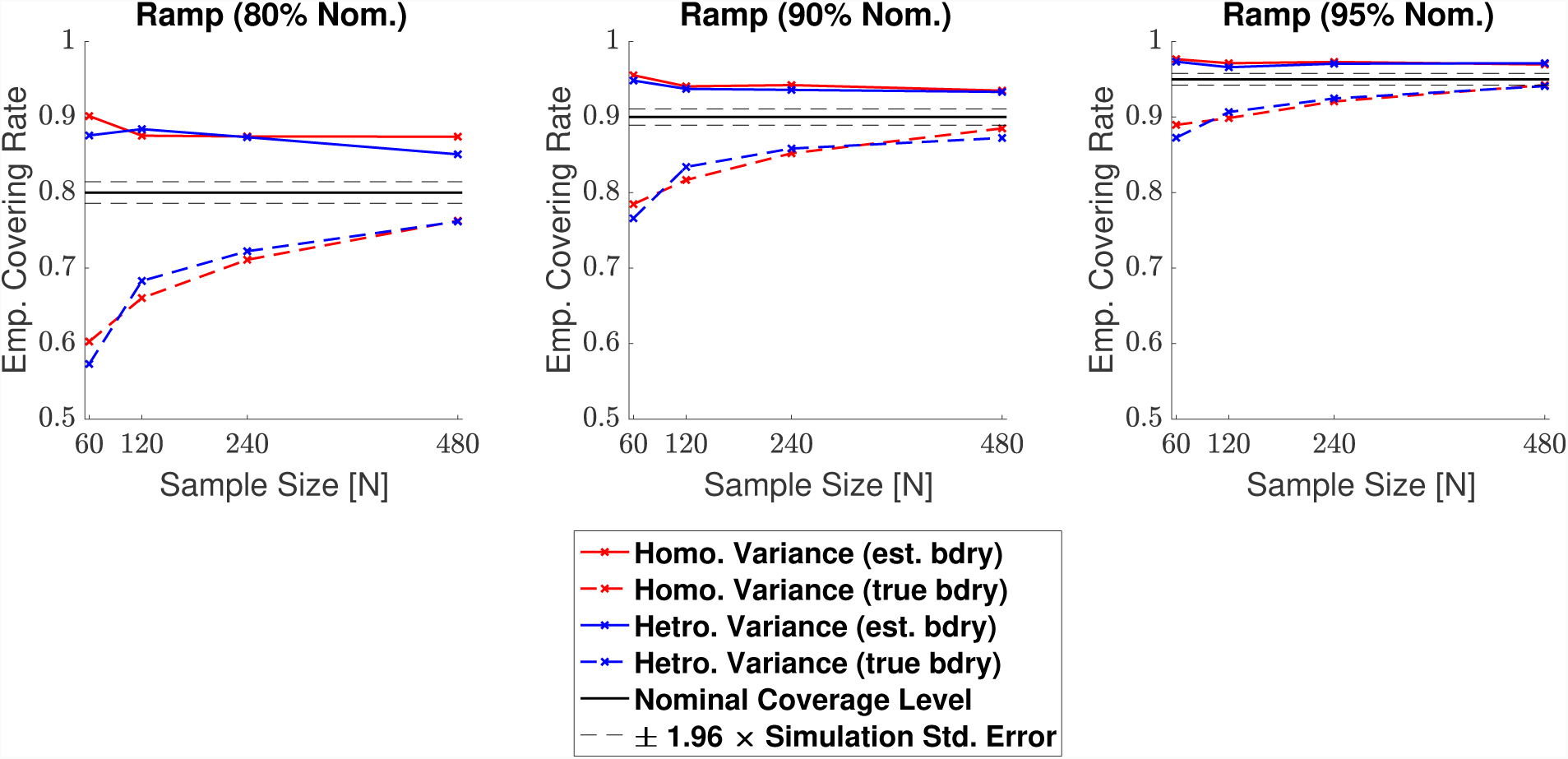
Coverage results for **Signal 1.**, the 2D linear ramp signal. While the true boundary coverage results (dashed curves) fall under the nominal level, results for the estimated boundary method (solid curves) that must be applied to real data remain above the nominal level. Performance of the method improved for larger confidence levels, and in particular, the estimated boundary results for a 95% confidence level seen in the right plot hover slightly above nominal coverage for all sample sizes.

**Figure 7.2:**
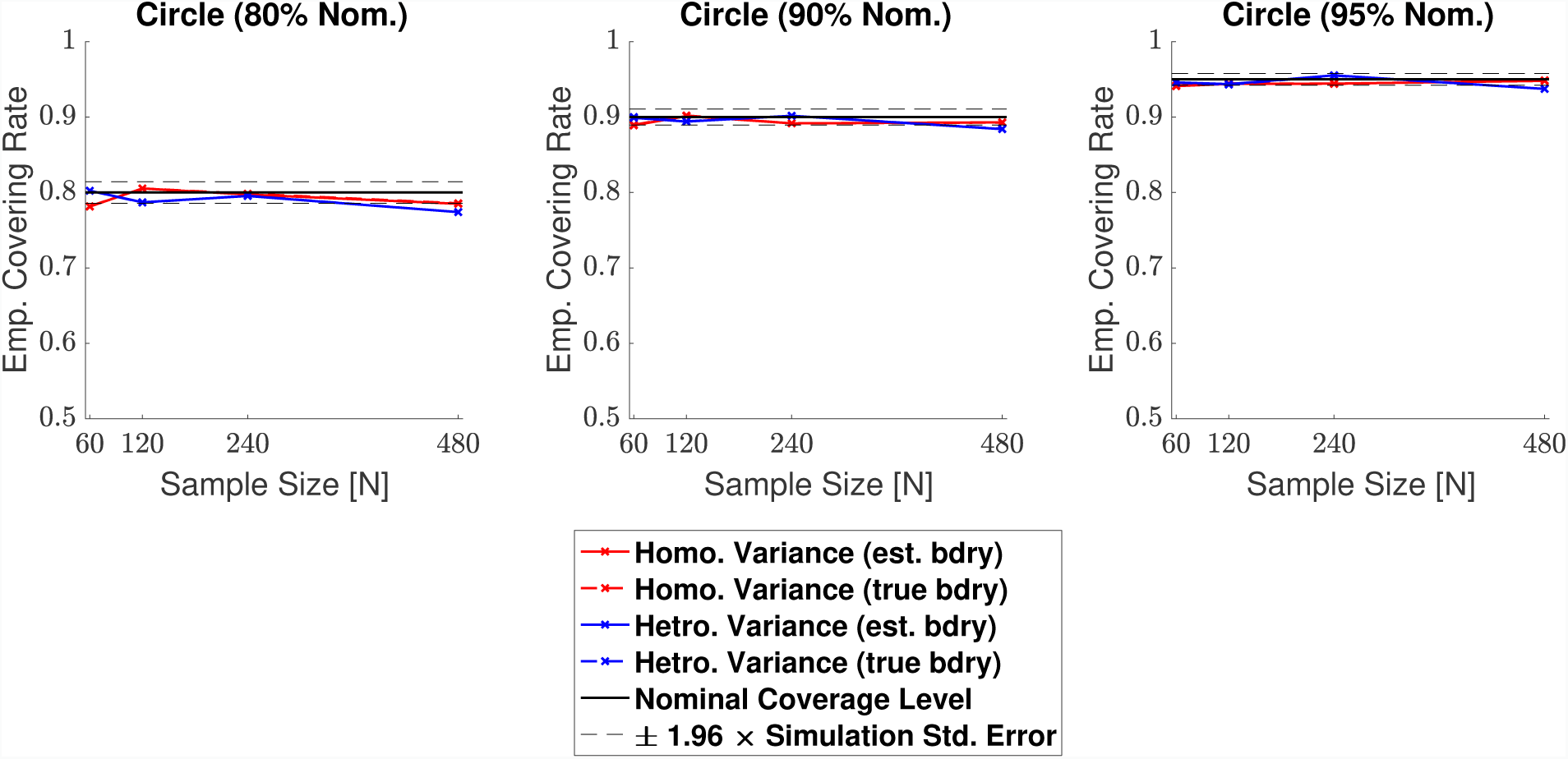
Coverage results for **Signal 2.**, the 2D circular signal. Coverage performance was close to nominal level in all simulations. The method was robust as to whether the subject-level noise had homogeneous (red curves) or heterogeneous variance (blue curves), or as to whether the estimated boundary (dashed curves) or true boundary (solid curves) method was used; in all plots, all of the curves lie practically on top of each other.

For the linear ramp, across all confidence levels we observed valid, over-coverage for the estimated boundary method, and under-coverage for the true boundary method. In both cases, the degree of agreement between our empirical results and the nominal coverage level improved for larger confidence levels, and as the sample size increased. For instance, while our estimated boundary empirical results were around 88% when the nominal target level was set at 80% (Fig. 7.1, left), corresponding empirical coverage results hovered around 97% for a nominal target of 95% (Fig. 7.1, right). Comparing the differences between the solid and dashed curves, there is also greater harmonization between the estimated and true boundary results for higher confidence levels. The method performed similarly regardless of whether homogeneous or heterogeneous noise was added to the model, evidenced by the minimal differences between the red and the blue curves for each of the two boundary methods seen in the plots.

For the circular signal the method performed remarkably well, with almost all our empirical coverage results lying within the 95% confidence interval of the nominal coverage rate (red and blue curves sandwiched between black dashed lines for all three plots in Fig. 7.2). Once again, the use of homogeneous or heterogeneous noise in the model had minimal difference on the method’s empirical coverage performance, and in this setting, our results were virtually identical whether the estimated boundary or true boundary was used for the bootstrap procedure. This has made the dashed curves hard to distinguish in the plots, as the solid curves lie practically on top of them.

### 4.3. 3D Simulations

Empirical coverage results for each of the three confidence levels 1 − α = 0.80, 0.90 and 0.95, are presented in Figs. 8.1, 8.2, 8.3 and 8.4 respectively for each of the four signal types (small sphere, large sphere, multiple spheres, Biobank full mean) displayed in Fig. 5. Results are also presented in tabular format in Table. S2. Once again, results obtained for simulations applying the bootstrap procedure over the estimated boundary 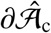 are displayed with a solid line, and results for simulations using the true boundary 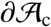 are displayed with a dashed line.

**Figure 8.1:**
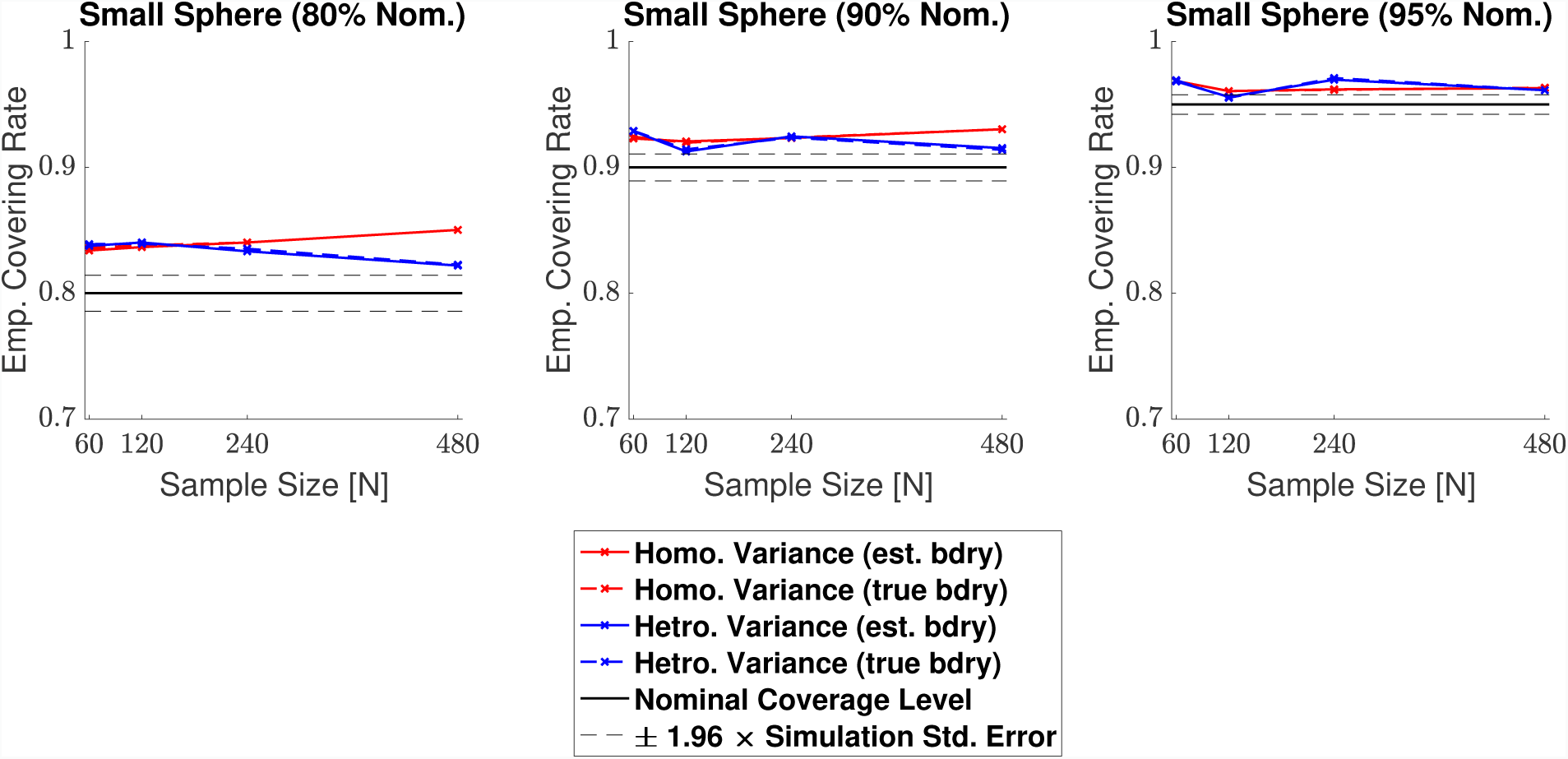
Coverage results for **Signal 1.**, the 3D small spherical signal. For all confidence levels, coverage remained above the nominal level in all simulations, and for a 95% confidence level (right plot), coverage hovered slightly above the nominal level for all sample sizes. The method was robust as to whether the subject-level noise had homogeneous (red curves) or heterogeneous variance (blue curves), or as to whether the estimated boundary (dashed curves) or true boundary (solid curves) method was used.

**Figure 8.2:**
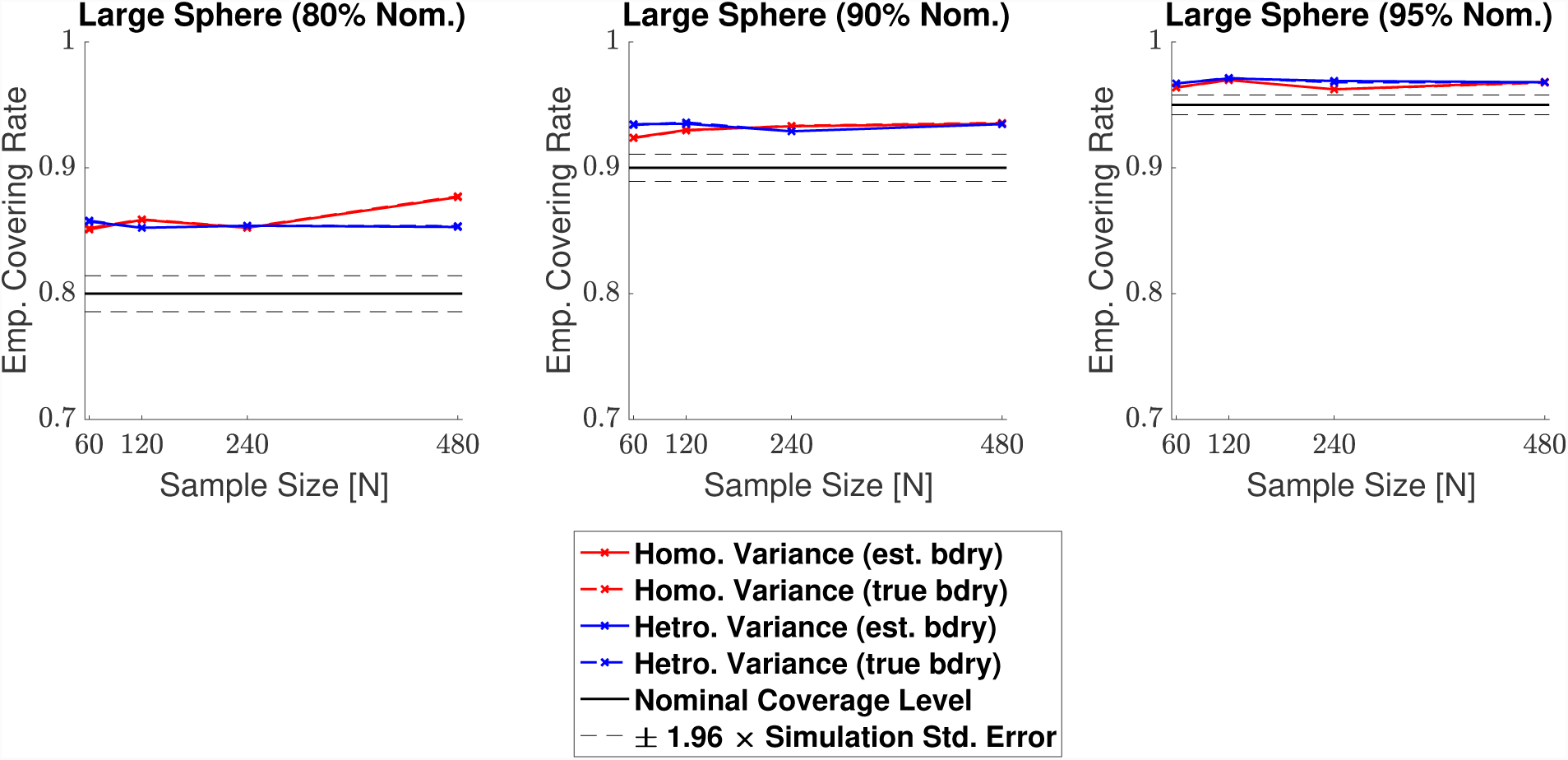
Coverage results for **Signal 2.**, the large 3D spherical signal. Coverage results here were very similar to the results for the small spherical signal shown in Fig. 8.1, suggesting that the method is robust to changes in boundary length.

**Figure 8.3:**
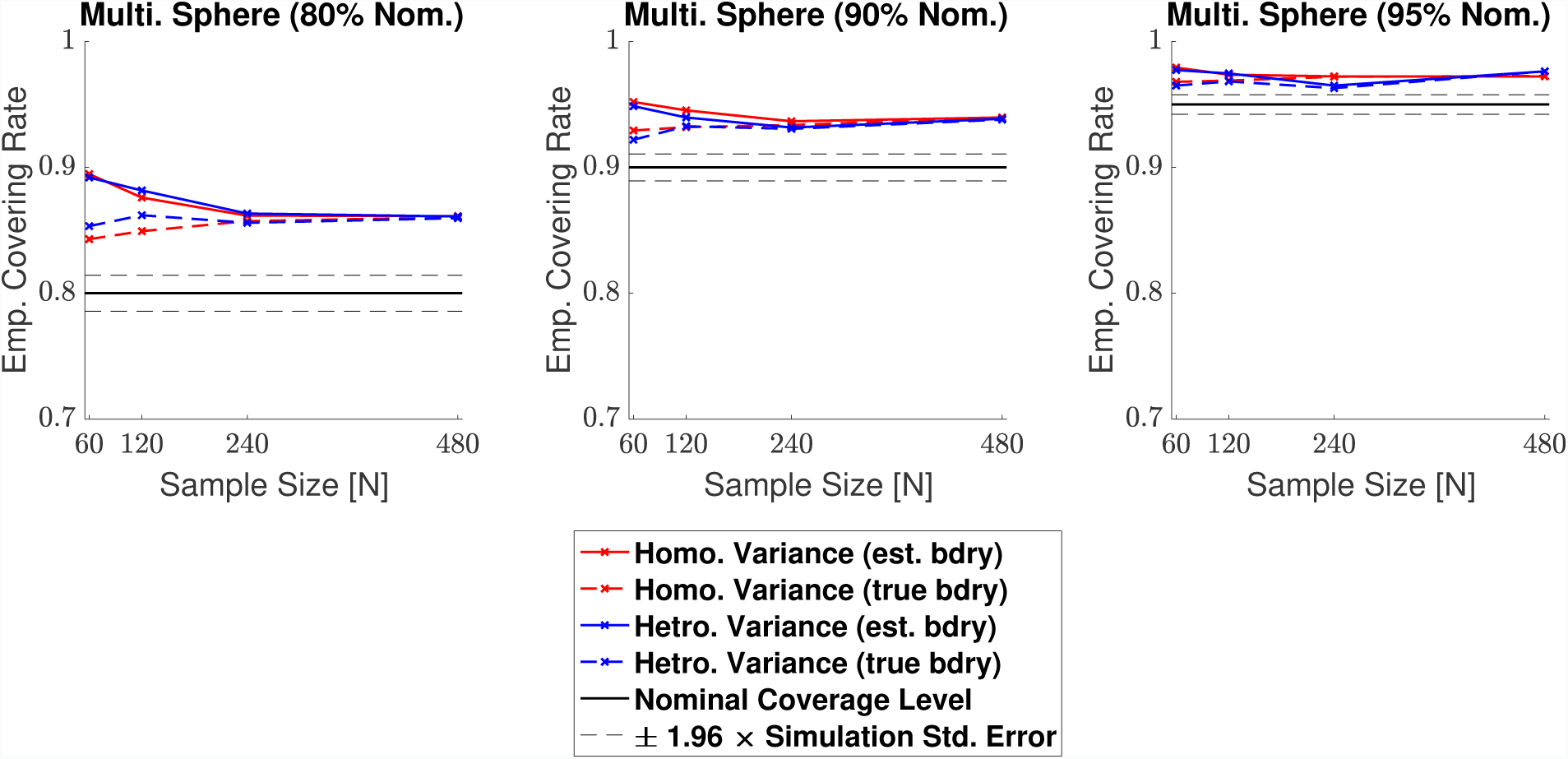
Coverage results for **Signal 3.**, the multiple spheres signal. Once again, for all confidence levels, coverage remained above the nominal level in all simulations. Here, the true boundary method (dashed curves) performed slightly better than the estimated boundary method (solid curves) in small sample sizes, although the choice of boundary made less of a difference for a higher confidence level. For a 95% confidence level (right plot), all results hover slightly above nominal coverage for all sample sizes.

**Figure 8.4:**
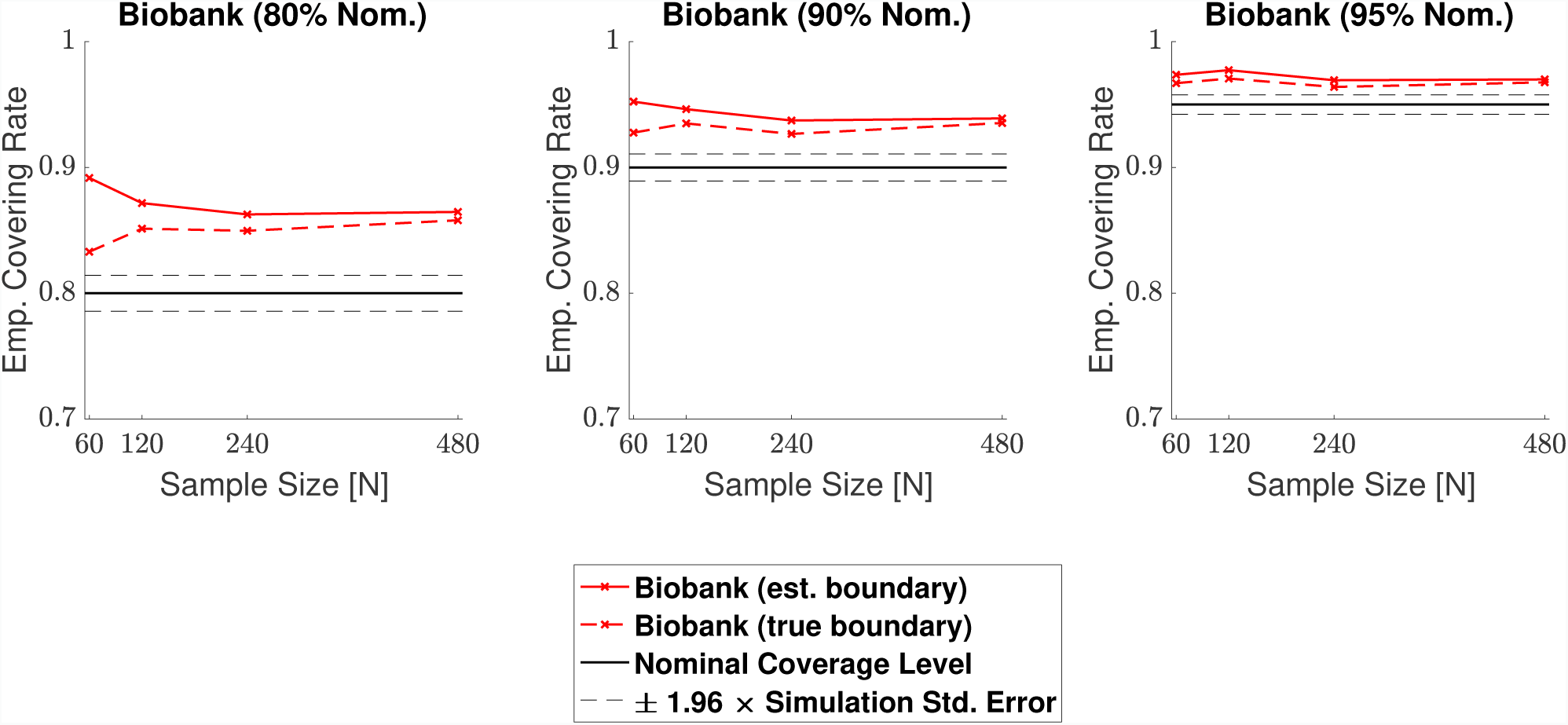
Coverage results for **Signal 4.**, the UK Biobank full mean signal, where the full standard deviation image was used as the standard deviation of the subject-level noise fields. Coverage results here were similar to the results for the multiple spheres signal shown in Fig. 8.3: In small sample sizes, coverage was slightly improved for the true boundary method (dashed curves) compared to the estimated boundary method (solid curves), however, for a 95% confidence level (right plots), all results hover slightly above nominal coverage for all sample sizes.

Overall, the results for all four signal types were consistent: In general, empirical coverage always came above the nominal target level, and the extent of over-coverage diminished when a higher confidence level was used. Particularly, for a nominal target of 1 − α = 0.95, all of our 3D empirical coverage results lie between 95% and 98%. The method was robust as to whether the bootstrap procedure was applied over the true or estimated boundary, or as to whether the variance of the noise field was homo- or heterogeneous. The similarity of the empirical coverage results, in spite of differences in these specific settings, is exhibited in all of the plots by the uniformity of the red and blues curves (indicating minimal differences in performance whether the noise had homogeneous or heterogeneous variance), and agreement between the solid and dashed curves (indicating minimal differences in performance whether the true boundary or estimated boundary was used). In the empirical coverage plots for the small and large spherical signals shown in Figs. 8.1 and 8.2, all of these curves lie virtually on top of each other.

While performance with the multiple spheres and Biobank signals presented in Figs. 8.3 and 8.4 was slightly better when using the true boundary, the true- and estimated boundary performance converged as the sample size increased.

### 4.4. Human Connectome Project

Confidence Sets obtained from applying the method to 80 subjects contrast data from the Human Connectome Project working memory task are shown in Fig. 9 and Fig. 10.

**Figure 9:**
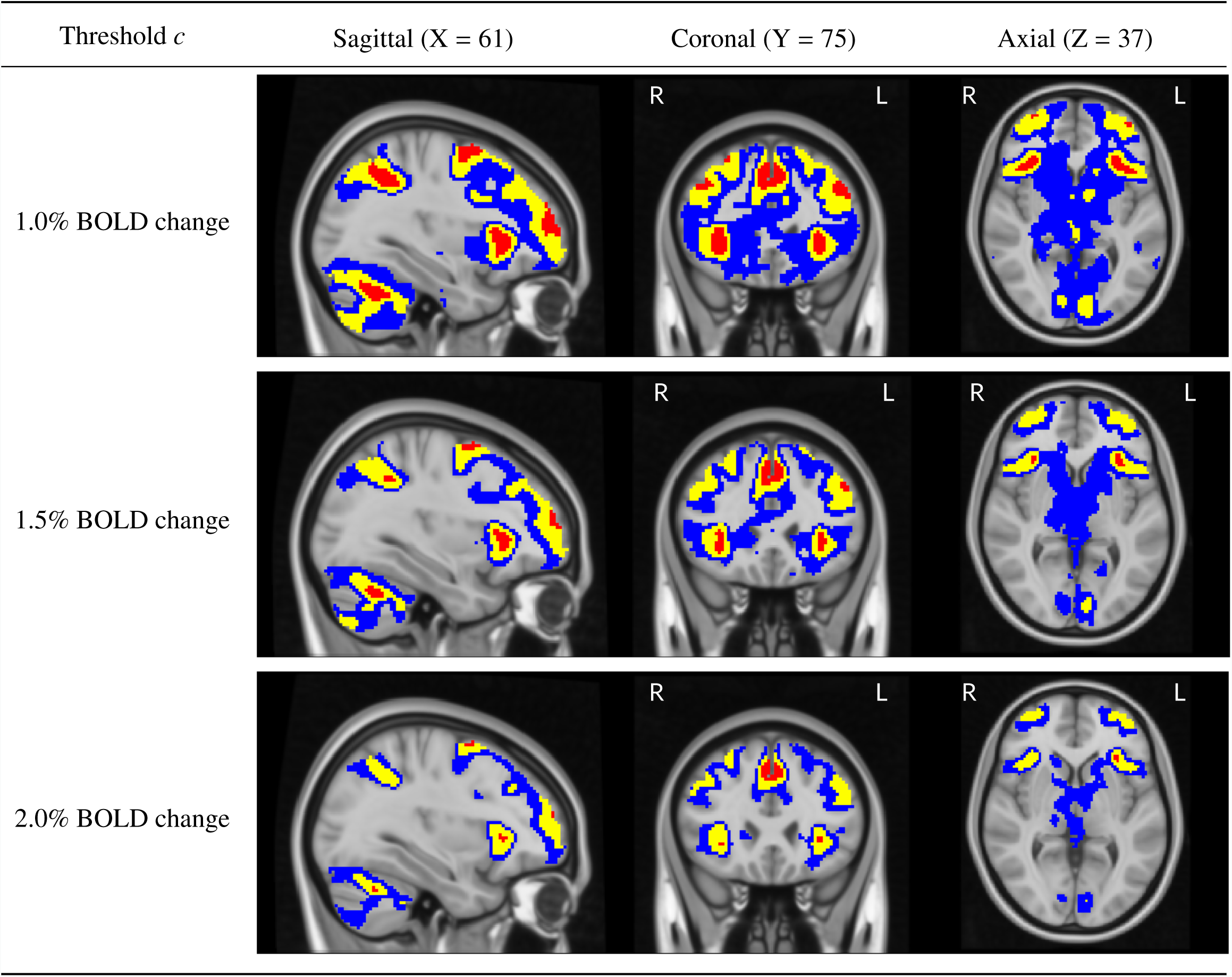
Slice views of the Confidence Sets for 80 subjects data from the HCP working memory task for *c* = 1.0%, 1.5% and 2.0% BOLD change thresholds. The upper CS 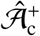 is displayed in red, and the lower CS 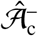 displayed in blue. In yellow is the point estimate set 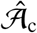, the best guess from the data of voxels that surpassed the BOLD change threshold. The red upper CS has localized regions in the frontal gyrus, frontal pole, anterior insula, supramarginal gyrus and cerebellum for which we can assert with 95% confidence that there has been (at least) a 1.0% BOLD change raw effect.

**Figure 10:**
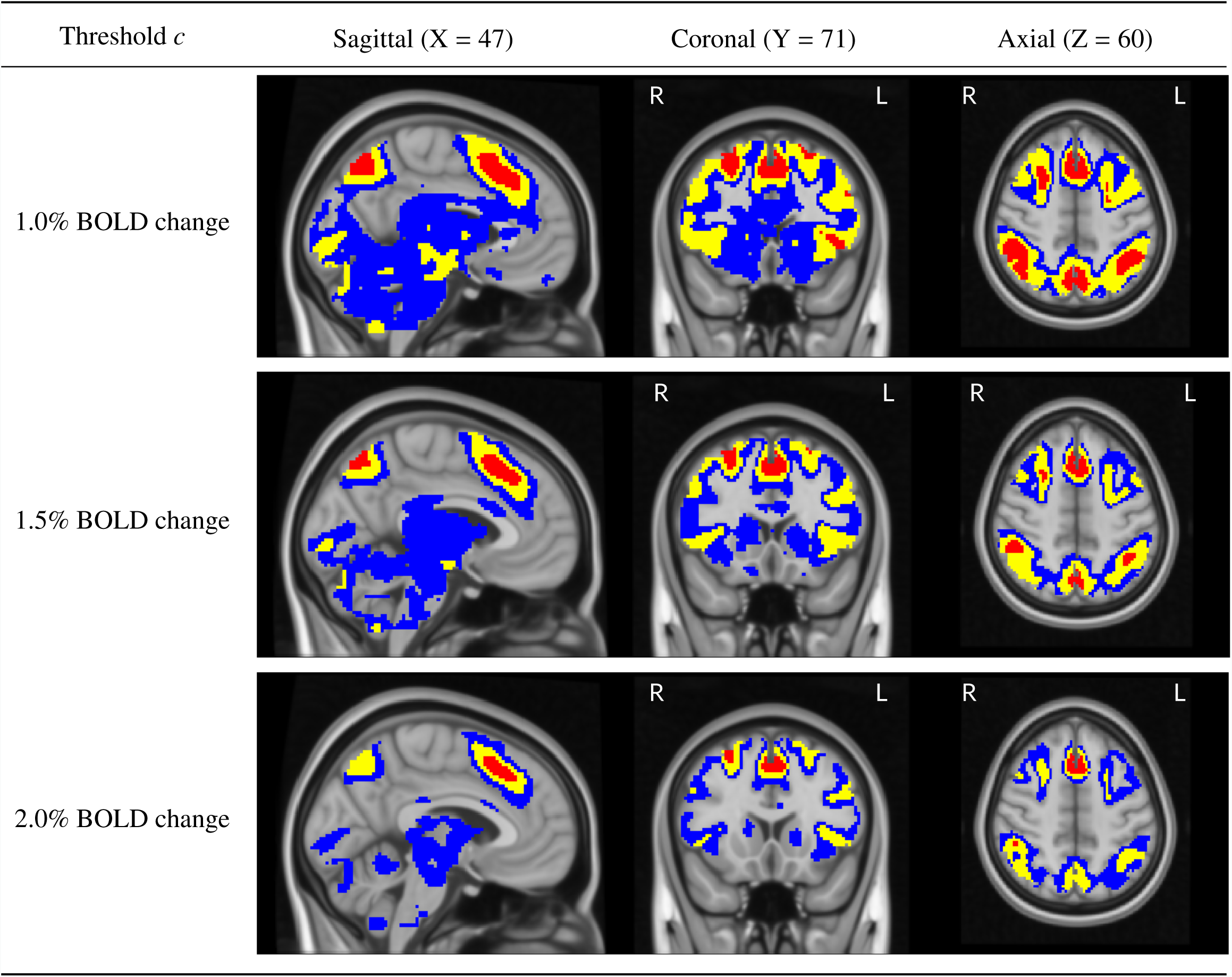
Further slice views of the Confidence Sets. Here, we see that the red upper CS has also localized regions in the anterior cingulate, superior front gyrus, supramarginal gyrus, and precuneous for which we can assert with 95% confidence that there has been (at least) a 1.0% BOLD change raw effect.

In both Fig. 9 and Fig. 10, the red upper CS localized brain regions within the frontal cortex commonly associated to working memory. This included areas of the middle frontal gyrus (left and right; Fig. 9, sagittal and coronal slices), superior frontal gyrus (left and right, Fig. 10, coronal slice) anterior insula (left and right; Fig. 9, sagittal and axial slices), as well as the anterior cingulate (Fig. 10, all slices). In all of the above regions, the method identified clusters of voxels for which we can assert with 95% confidence there was a percentage BOLD change raw effect greater than 2.0% (Fig. 9 and Fig. 10, bottom plots).

Further brain areas localized by the upper CS were the frontal pole (left and right; Fig. 9, sagittal and axial slices), supramarginal gyrus (left and right; Fig. 9, sagittal slice and Fig. 10, coronal and axial slices), precuneous (Fig. 10, sagittal slice) and cerebellum (Fig. 9, sagittal slice). While for these areas we can assert with 95% confidence there was a percentage BOLD change raw effect greater than at least 1.0% (Fig. 9 and Fig. 10, top plots), on-the-whole the method only localized areas where there was a BOLD change of at least 2.0% in parts of the frontal cortex. This can be observed by the ‘disappearance’ of the red CSs in brain regions located in the ocipital lobe for the 2.0% BOLD change plots when compared with the corresponding 1.0% and 1.5% BOLD change plots in Fig. 9 and Fig. 10.

As the percentage BOLD change threshold increases between plots, there is a shrinking of both the blue lower CSs and red upper CSs: By using a larger threshold, there are less voxels we can confidently declare have surpassed this higher level of percentage BOLD change, and thus the volume of the red upper CSs decreases (in some cases, vanishing). At the same time, there are more voxels we expect to be able to confidently declare have fallen below the threshold. Since these are precisely the (grey background) voxels that lie outside of the lower blue CSs, the volume of the blue lower CSs also decreases.

Finally, in Fig. S1 and Fig. S2 the red upper CSs are compared with the thresholded *t*-statistic map (green-yellow voxels) obtained from applying a traditional one-sample *t*-test group-analysis to the 80 subjects working memory task contrast data, using a voxelwise FWE-corrected threshold of *p* < 0.05. Differences here highlight how statistical significance may not translate to practical significance; while over 28,000 voxels were declared as active in the thresholded *t*-statistic results, only 4,818 voxels were contained in the upper CS indicating a percentage BOLD change of at least 1.0%.

## 5. Discussion

### 5.1. Spatial Inference on %BOLD Raw Effect Size

Thorough interpretation of neuroimaging results requires an appreciation of the practical (as well as statistical) significance of differences through visualization of raw effect magnitude maps with meaningful units (Chen et al., 2017). In this work, we have presented a method to create confidence sets for raw effect size maps, providing formal confidence statements on regions of the brain where the %BOLD response magnitude has exceeded a specified activation threshold, alongside regions where the %BOLD response has *not* surpassed this threshold. Both of these statements are made simultaneously, and across the entire brain. This not only enables researchers to infer brain areas that have responded to a task, but also allows for inference on areas that did not respond to the task. In this sense, the method goes beyond statistical hypothesis testing, where the null-hypothesis of no activation can ‘fail to be rejected’, but never accepted. By operating on percentage BOLD change units, instead of *t*-statistic values, the confidence set maps present a clear and more direct interpretation of the biophysical changes that occur during a neuroimaging study, which can be distorted by the thresholded statistic maps commonly reported at the end of an investigation (Engel and Burton, 2013). In essence, the CSs synthesize information that is usually provided separately in a raw effect size and *t*-statistic map, determining practically significant effects in terms of effect magnitude, that are also reliable in terms of statistical significance traditionally given by *p*-values in a statistic image. While in this work we have focused on BOLD fMRI, the methods presented here are applicable to any neuroimaging measure that can be fit in a group-level GLM.

The choice of threshold *c* is ultimately up to the user, and may depend on the aims of the investigation. Researchers may choose a threshold based on prior knowledge of raw effect sizes observed in previous similar studies, and it is likely that localization of larger raw effects will be possible as sample sizes increase. Obtaining the CSs for the Human Connectome Project contrast data in this work was computationally quick, each analysis taking no longer than a couple of minutes. Therefore, one possible strategy is to evaluate a variety of different *c*’s on pilot or historical data before fixing a value to use on a study of interest.

### 5.2. Analysis of HCP data and Simulation Results

In our analysis of the HCP emotional faces task-fMRI dataset, we have primarily focused on activated areas localized by the red upper CS. However, the confidence set maps in Fig. 9 and Fig. 10 also quantify the spatial precision of the point estimate ‘best guess from the data’ activation clusters. While so far we have described the confidence sets in terms of the red and blue upper and lower CSs, we now highlight that the set difference between the upper and lowers CSs acts as a confidence region itself; with 95% confidence, we can assert that the boundary of the point estimate set (raw effect size > threshold; yellow voxels overlapped by red in Fig. 9 and Fig. 10) is completely contained within this region. The set difference region, visualized by blue and yellow voxels (but not red) in Fig. 9 and Fig. 10, therefore anticipates how the point estimate clusters may fluctuate if the experiment was to be repeated again. From this perspective, the vast areas of the brain covered by blue in Fig. 9 and Fig. 10 demonstrate the high level of uncertainty in localizing a raw effect size of, for example, 1.0% BOLD change, despite the large sample size of *N* = 80 used for the HCP. The regions of greatest uncertainty were sub-cortical areas, covered by expansive clusters of blue as seen in the axial slices displayed in Fig. 9 and sagittal slices in Fig. 10. Large intersubject variability here may be explained by the high multi-band acceleration factor used in the HCP scanning protocol, which is generally more suited for scanning the cortex (Smith et al., 2013).

For the 2D simulations, the method achieved close to nominal coverage for the circular signal, but performed less well for the ramp signal, obtaining under-coverage for the true boundary method and over-coverage for the estimated boundary method. We believe differences in the circle and ramp results are not due to changes in the signal shape per se, but instead are caused by differences in the slope of each shape close to the true boundary 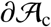. Since the linear ramp signal has a shallower gradient at the true boundary compared to the circle, local changes in the observed signal around the boundary are dominated by changes in the noise. Since the noise is more wavey than the signal, the linear interpolation method for obtaining the boundary is likely to be less accurate for the ramp, causing too many violations of the subset condition, which may explain the under-coverage for the true boundary results seen here.

For the 3D simulations, the method obtained over-coverage in all of our results. Here, the degree of over-coverage was consistently larger for the smaller confidence level of 1 − α = 0.80 in comparison to the larger confidence level of 1 − α = 0.95. Notably, the over-coverage was also more severe for signals with a longer boundary, such as the multiple spheres and Biobank signals, when compared to the Small Sphere signal that had a shorter boundary length. One possible reason for this is that our proposed method for assessing coverage may still be missing instances where violations of the subset condition 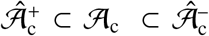 occur, causing the results to be slightly positively biased. While our assessment method reduces the influence of grid coarseness by sampling locations on the true continuous boundary 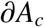, ultimately we can still only assess coverage at a discrete set of points on a continuous process. For signals with a longer boundary length, the set of sampled locations obtained with the interpolation method is relatively less dense within the true continuous boundary, and thus it is more likely violations of the subset condition are missed. Over-coverage for smaller confidence levels may also be explained by this, as theoretically more violations should occur here, but these may be missed due to inaccuracies caused by the discreteness of the lattice. This line of reasoning is consistent with Section 4.4 of *SSS*, where it was shown that coverage approached the nominal level as the resolution of the grid was increased.

### 5.3. Methodological Innovations

In this work, we have advanced on the original methods applied in *SSS*. From a theoretical standpoint, we have proposed a Wild *t*-Bootstrap method (dividing bootstrap residuals by bootstrap standard deviation) to compute the critical quantile value *k*. We have also introduced an interpolation scheme for obtaining the boundary and assessing the simulation coverage results to reduce the influence of grid coarseness. In Section 4.1, we demonstrated that applying the assessment method in *SSS* could lead to empirical coverage of close to 100%, suggesting that this method may considerably bias the simulation results upwards. When using our proposed assessment, the Wild Bootstrap method suffered from under-coverage, most severely for small sample sizes in the 3D setting of the large spherical signal presented in Fig. 6.2. This was greatly remedied by the Wild *t*-Bootstrap method, for which empirical results stayed close to the nominal target independent of sample size.

Our simulations using the original procedures may not seem consistent with the simulation results published in Figure 5 of *SSS*, where empirical coverage stayed close to the nominal target. However, the signal-plus-noise models investigated to test the performance of the CSs in *SSS* were much smoother than the synthetic signals considered to emulate fMRI data with this effort. By applying a larger degree of smoothing, the signals used in *SSS* effectively had a much higher resolution. Because of this, it is likely the resolution issue presented in Fig. 3 was less critical, reducing the positive bias in empirical coverage induced from using the original simulation assessment procedure. Further evidence for this is provided in Figure 7 of *SSS*, where they observed an increase in coverage after repeating their simulations on a coarser lattice. In our simulation results in Section 4.1, the scale of under-coverage from using the Gaussian Wild bootstrap method was much more severe for the 3D simulation on the spherical signal in Fig. 6.2 compared to the 2D circular signal in Fig. 6.1. This may explain why the Gaussian Wild bootstrap method performed relatively well in *SSS*, as only 2D signals were considered there.

### 5.4. Limitations & Future Work

The principal limitation of this work is one that is intentional and explicit: Our method is for spatial inference on maps of raw and not standardized effects, such as Cohen’s *d* or partial *R*^2^ (*t*or *F*-statistics, which scale with sample size, do not estimate population quantities and are not suitable for making confidence statements). Even when scaled to percentage BOLD change, it has been shown that raw effects can modulate with acquisition parameters such as the scanner field strength or echo time (UIudag et al., 2009). Users should therefore be cautious when combining effect estimates from studies using heterogeneous acquisition setups, and clearly specify such differences when reporting the results of any meta-analysis on raw effects. It is also known that inhomogeneities in the vasculature of the brain is a cause of variation in the BOLD response. Therefore, we recommend that any interpretation of %BOLD change inferred from the CSs is referenced against a variance map or similar image that indicates the most venous brain regions. We note that each of these points are general complications of raw effect sizes within fMRI, rather than issues with the method proposed in this effort per se. Nonetheless, the use of standardized effect estimates may help to remedy these problems in the future. The statistical characteristics of standardized effect maps are fundamentally different to the raw effect images motivating the method here, and the topic of our current work is to develop CSs for standardized effect size images.

The need for resampling to conduct inference is another limitation of this effort, especially given the big data motivation of this work. However, the bootstrap is only conducted on the estimated boundary, 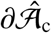, not the whole 3D volume, which substantially reduces the computational burden. For very large datasets, techniques for approximating empirical distributions can be used to improve the accuracy of the estimation of *k* based on a smaller number (e.g. *B* = 500) of bootstrap samples (Winkler et al., 2016).

## 6. Contributions

A.B. drafted the manuscript. All authors edited and revised the manuscript. A.B. proposed use of Rademacher variables for the bootstrap and developed the method for assessing simulations. F.T. proposed use of the Wild *t*-Bootstrap method and developed the linear interpolation method for approximating the boundary. A.B. conducted simulations and data analysis, with additional contributions from F.T. to the simulation and simulation figures code. A.S. and T.E.N. oversaw the project in all its intellectual, methodological and computational aspects.

## 7. Data Availability

We have used data from The Human Connectome Project and UK Biobank. All code used for the simulations and analysis of HCP data are available at: https://github.com/AlexBowring/Spatial_Confidence_Sets_Raw.

## 8. Acknowledgements

TEN is supported by the Wellcome Trust (100309/Z/12/Z). AS and FT were partially supported by NIH grants (R01CA157528 and R01EB026859). The authors would like to thank Max Sommerfeld for his early involvement in the project, and Samuel Davenport for helping to process the Biobank Data used in the Simulations section.

## Footnotes

1 For examples of how to set up more complex designs and contrasts, see Figure A.2. in the Appendix A section of (Poldrack et al., 2011).

## Appendix A. Supplementary Human Connectome Project Results

**Supplementary Figure 1:**
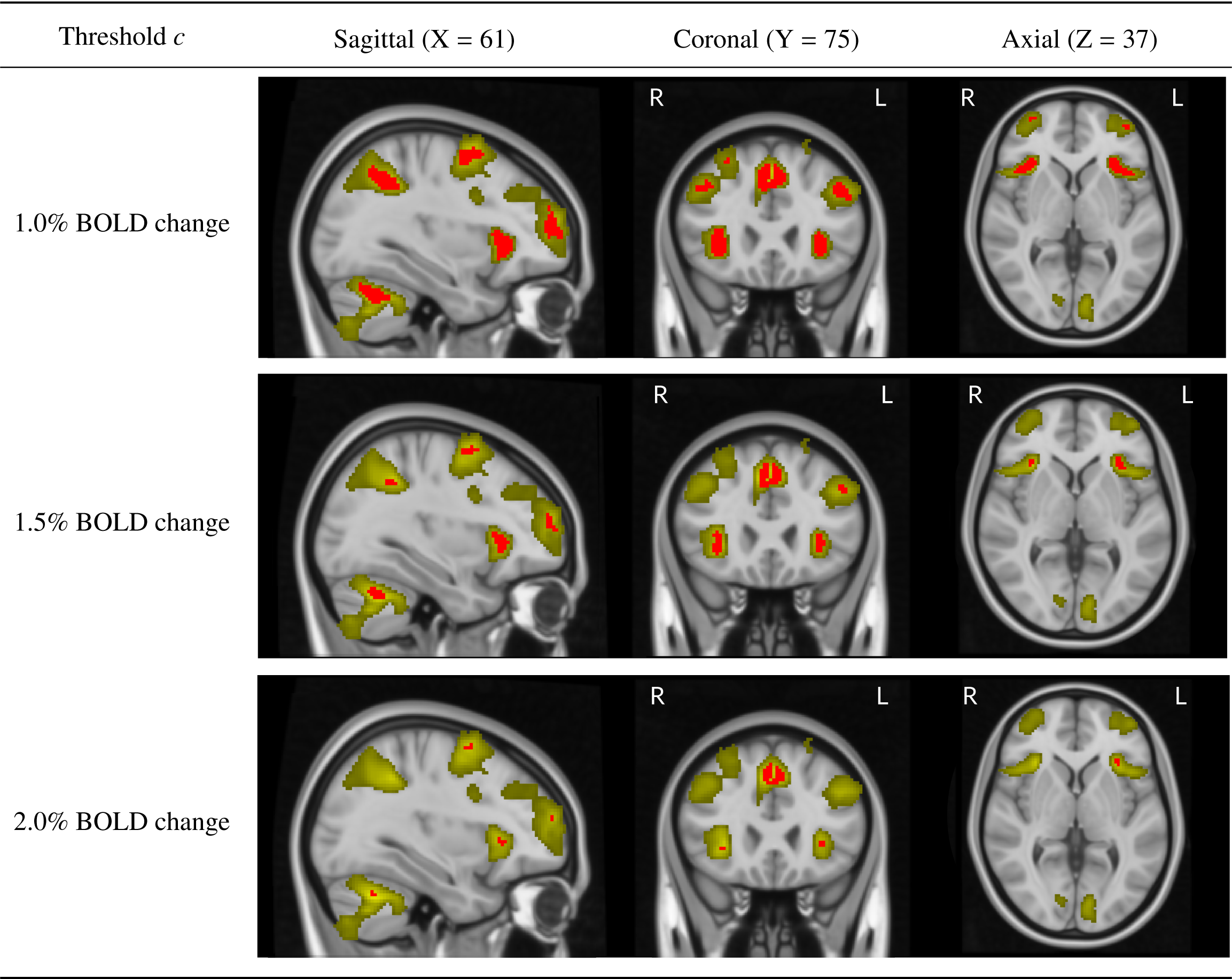
Comparing the upper Confidences Sets for the HCP working memory task data (same slice views as Fig. 9) with the thresholded *t*-statistic results obtained by applying a traditional group-level one-sample *t*-test, voxelwise *p* < 0.05 FWE correction (green-yellow voxels). While over 25,000 voxels were determined as statistically significant with the standard inference method, less than 5,000 voxels were asserted to have at least a 1.0% BOLD change by the CSs. In particular, the two statistically significant clusters spanning the left and right side of the frontal lobe contained almost no voxels with a practical effect size of over 1.5% BOLD change.

**Supplementary Figure 2:**
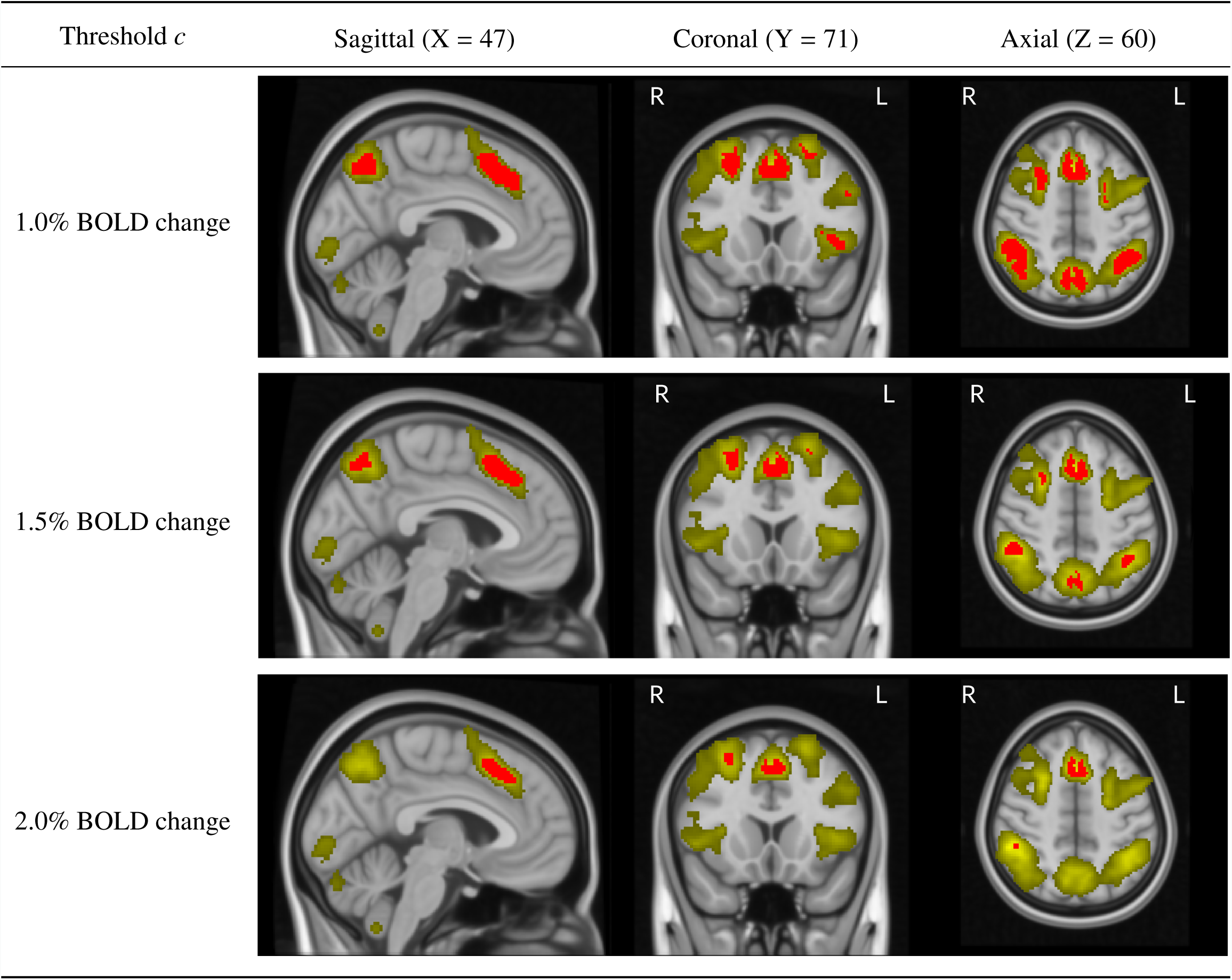
Comparing the upper Confidences Sets for the HCP working memory task data (same slice views as Fig. 10) with the thresholded *t*-statistic results obtained by applying a traditional group-level one-sample *t*-test, voxelwise *p* < 0.05 FWE correction (green-yellow voxels). While one large statistically significant cluster covers the supramarginal gyrus, angular gyrus and precuneous, the CSs localize the precise areas with practically significant effect sizes within each of these regions.

**Supplementary Table 1.** Empirical coverage results for the 2D simulations using nominal (nom.) coverage levels 1 − α = 80%, 90% and 95%. Results are shown for applying the Wild *t*-Bootstrap method to the residual field along the estimated boundary 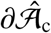 (top) and the true boundary 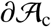(bottom).

|  | 2D Signal 1. (Ramp) |  | 2D Signal 2. (Circle) |  |
| --- | --- | --- | --- | --- |
|  | Standard Dev 1. | Standard Dev 2. | Standard Dev 1. | Standard Dev 2. |
| $\partial\hat{\mathcal{A}}_c$ | | | | |
| 80% nom. |  |  |  |  |
| $N = 60$ | 90.13% ± 0.54% | 87.57% ± 0.60% | 78.13% ± 0.75% | 80.23% ± 0.73% |
| 120 | 87.53% ± 0.60% | 88.40% ± 0.58% | 80.53% ± 0.72% | 78.70% ± 0.75% |
| 240 | 87.43% ± 0.61% | 87.33% ± 0.61% | 79.73% ± 0.73% | 79.53% ± 0.74% |
| 480 | 87.40% ± 0.61% | 85.07% ± 0.65% | 78.50% ± 0.75% | 77.40% ± 0.76% |
| 90% nom. |  |  |  |  |
| $N = 60$ | 95.53% ± 0.38% | 94.83% ± 0.40% | 88.90% ± 0.57% | 89.90% ± 0.55% |
| 120 | 94.07% ± 0.43% | 93.73% ± 0.44% | 90.13% ± 0.54% | 89.40% ± 0.56% |
| 240 | 94.23% ± 0.43% | 93.60% ± 0.45% | 89.17% ± 0.57% | 90.17% ± 0.54% |
| 480 | 93.50% ± 0.45% | 93.33% ± 0.46% | 89.30% ± 0.56% | 88.40% ± 0.58% |
| 95% nom. |  |  |  |  |
| $N = 60$ | 97.67% ± 0.28% | 97.33% ± 0.29% | 94.10% ± 0.43% | 94.60% ± 0.41% |
| 120 | 97.13% ± 0.30% | 96.60% ± 0.33% | 94.40% ± 0.42% | 94.37% ± 0.42% |
| 240 | 97.30% ± 0.30% | 97.07% ± 0.31% | 94.43% ± 0.42% | 95.53% ± 0.38% |
| 480 | 96.97% ± 0.31% | 97.13% ± 0.30% | 94.80% ± 0.41% | 93.73% ± 0.44% |
| $\partial\mathcal{A}_c$ | | | | |
| 80% nom. |  |  |  |  |
| $N = 60$ | 60.27% ± 0.89% | 57.30% ± 0.90% | 78.17% ± 0.75% | 80.23% ± 0.73% |
| 120 | 66.03% ± 0.86% | 68.30% ± 0.85% | 80.53% ± 0.72% | 78.67% ± 0.75% |
| 240 | 71.10% ± 0.83% | 72.23% ± 0.82% | 79.83% ± 0.73% | 79.57% ± 0.74% |
| 480 | 76.27% ± 0.78% | 76.17% ± 0.78% | 78.57% ± 0.75% | 77.40% ± 0.76% |
| 90% nom. |  |  |  |  |
| $N = 60$ | 78.47% ± 0.75% | 76.60% ± 0.77% | 88.97% ± 0.57% | 90.00% ± 0.55% |
| 120 | 81.67% ± 0.71% | 83.40% ± 0.68% | 90.20% ± 0.54% | 89.43% ± 0.56% |
| 240 | 85.20% ± 0.65% | 85.83% ± 0.64% | 89.17% ± 0.57% | 90.17% ± 0.54% |
| 480 | 88.50% ± 0.58% | 87.23% ± 0.61% | 89.27% ± 0.57% | 88.43% ± 0.58% |
| 95% nom. |  |  |  |  |
| $N = 60$ | 88.97% ± 0.57% | 87.27% ± 0.61% | 94.17% ± 0.43% | 94.57% ± 0.41% |
| 120 | 89.87% ± 0.55% | 90.67% ± 0.53% | 94.47% ± 0.42% | 94.30% ± 0.42% |
| 240 | 92.07% ± 0.49% | 92.47% ± 0.48% | 94.40% ± 0.42% | 95.50% ± 0.39% |
| 480 | 94.23% ± 0.43% | 94.10% ± 0.43% | 94.87% ± 0.40% | 93.73% ± 0.44% |

**Supplementary Table 2.** Empirical coverage results for the 3D simulations using nominal (nom.) coverage levels 1 − α = 80%, 90% and 95%. Results are shown for applying the Wild *t*-Bootstrap method to the residual field along the estimated boundary 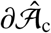(top) and the true boundary 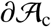(bottom).

|  | 3D Signal 1. (Small Sphere) |  | 3D Signal 2. (Large Sphere) |  |
| --- | --- | --- | --- | --- |
|  | Standard Dev 1. | Standard Dev 2. | Standard Dev 1. | Standard Dev 2. |
| $\partial\hat{\mathcal{A}}_c$ | | | | |
| 80% nom. |  |  |  |  |
| $N = 60$ | 83.40% ± 0.68% | 83.77% ± 0.67% | 85.10% ± 0.65% | 85.73% ± 0.64% |
| 120 | 83.67% ± 0.67% | 84.03% ± 0.67% | 85.87% ± 0.64% | 85.23% ± 0.65% |
| 240 | 84.03% ± 0.67% | 83.77% ± 0.67% | 85.23% ± 0.65% | 85.40% ± 0.64% |
| 480 | 85.03% ± 0.65% | 82.20% ± 0.70% | 87.67% ± 0.60% | 85.30% ± 0.65% |
| 90% nom. |  |  |  |  |
| $N = 60$ | 92.30% ± 0.49% | 92.87% ± 0.47% | 92.40% ± 0.48% | 93.47% ± 0.45% |
| 120 | 92.07% ± 0.49% | 91.27% ± 0.52% | 93.00% ± 0.47% | 93.50% ± 0.45% |
| 240 | 92.33% ± 0.49% | 92.87% ± 0.47% | 93.30% ± 0.46% | 92.90% ± 0.47% |
| 480 | 93.03% ± 0.46% | 91.53% ± 0.51% | 93.50% ± 0.45% | 93.47% ± 0.45% |
| 95% nom. |  |  |  |  |
| $N = 60$ | 96.87% ± 0.32% | 96.83% ± 0.32% | 96.40% ± 0.34% | 96.70% ± 0.33% |
| 120 | 96.07% ± 0.35% | 95.60% ± 0.37% | 96.97% ± 0.31% | 97.10% ± 0.31% |
| 240 | 96.20% ± 0.35% | 96.83% ± 0.32% | 96.23% ± 0.35% | 96.90% ± 0.32% |
| 480 | 96.30% ± 0.34% | 96.13% ± 0.35% | 96.83% ± 0.32% | 93.80% ± 0.44% |
| $\partial\mathcal{A}_c$ | | | | |
| 80% nom. |  |  |  |  |
| $N = 60$ | 83.60% ± 0.68% | 83.90% ± 0.67% | 85.20% ± 0.65% | 85.80% ± 0.64% |
| 120 | 83.80% ± 0.67% | 83.93% ± 0.67% | 85.90% ± 0.64% | 85.23% ± 0.65% |
| 240 | 84.03% ± 0.67% | 83.90% ± 0.67% | 85.27% ± 0.65% | 85.40% ± 0.64% |
| 480 | 85.03% ± 0.65% | 82.27% ± 0.70% | 87.73% ± 0.60% | 85.37% ± 0.65% |
| 90% nom. |  |  |  |  |
| $N = 60$ | 92.43% ± 0.48% | 92.90% ± 0.47% | 92.37% ± 0.48% | 93.40% ± 0.45% |
| 120 | 91.97% ± 0.50% | 91.43% ± 0.51% | 92.97% ± 0.47% | 93.60% ± 0.45% |
| 240 | 92.37% ± 0.48% | 92.90% ± 0.47% | 93.33% ± 0.46% | 92.90% ± 0.47% |
| 480 | 93.03% ± 0.46% | 91.40% ± 0.51% | 93.57% ± 0.45% | 93.47% ± 0.45% |
| 95% nom. |  |  |  |  |
| $N = 60$ | 96.87% ± 0.32% | 96.93% ± 0.31% | 96.37% ± 0.34% | 96.70% ± 0.33% |
| 120 | 96.07% ± 0.35% | 95.53% ± 0.38% | 96.97% ± 0.31% | 97.13% ± 0.30% |
| 240 | 96.17% ± 0.35% | 96.93% ± 0.31% | 96.23% ± 0.35% | 96.80% ± 0.32% |
| 480 | 96.33% ± 0.34% | 96.13% ± 0.35% | 96.77% ± 0.32% | 96.80% ± 0.32% |

Supplementary Table 2. (continued)
|  | 3D Signal 3. (Multiple Spheres) |  | 3D Signal 4. (UK Biobank) |
| --- | --- | --- | --- |
|  | Standard Dev 1. | Standard Dev 2. | UK Biobank SD |
| $\partial\hat{\mathcal{A}}_c$ | | | |
| 80% nom. |  |  |  |
| $N = 60$ | 89.47% ± 0.56% | 89.20% ± 0.57% | 89.17% ± 0.57% |
| 120 | 87.60% ± 0.60% | 88.17% ± 0.59% | 87.17% ± 0.61% |
| 240 | 86.17% ± 0.63% | 86.33% ± 0.63% | 86.27% ± 0.63% |
| 480 | 86.13% ± 0.63% | 86.10% ± 0.63% | 87.67% ± 0.60% |
| 90% nom. |  |  |  |
| $N = 60$ | 95.20% ± 0.39% | 94.87% ± 0.40% | 95.23% ± 0.39% |
| 120 | 94.53% ± 0.42% | 93.97% ± 0.43% | 94.63% ± 0.41% |
| 240 | 93.67% ± 0.44% | 93.17% ± 0.46% | 93.73% ± 0.44% |
| 480 | 93.97% ± 0.43% | 93.87% ± 0.44% | 93.50% ± 0.45% |
| 95% nom. |  |  |  |
| $N = 60$ | 97.93% ± 0.26% | 97.73% ± 0.27% | 97.37% ± 0.29% |
| 120 | 97.37% ± 0.29% | 97.47% ± 0.29% | 97.73% ± 0.27% |
| 240 | 97.23% ± 0.30% | 96.50% ± 0.34% | 96.93% ± 0.31% |
| 480 | 97.23% ± 0.30% | 97.63% ± 0.28% | 96.83% ± 0.32% |
| $\partial\mathcal{A}_c$ | | | |
| 80% nom. |  |  |  |
| $N = 60$ | 84.30% ± 0.66% | 85.33% ± 0.65% | 83.30% ± 0.68% |
| 120 | 84.93% ± 0.65% | 86.20% ± 0.63% | 85.13% ± 0.65% |
| 240 | 85.73% ± 0.64% | 85.60% ± 0.64% | 84.97% ± 0.65% |
| 480 | 86.03% ± 0.63% | 85.97% ± 0.63% | 87.73% ± 0.60% |
| 90% nom. |  |  |  |
| $N = 60$ | 92.93% ± 0.47% | 92.20% ± 0.49% | 92.77% ± 0.47% |
| 120 | 93.20% ± 0.46% | 93.27% ± 0.46% | 93.50% ± 0.45% |
| 240 | 93.37% ± 0.45% | 93.07% ± 0.46% | 92.67% ± 0.48% |
| 480 | 93.97% ± 0.43% | 93.80% ± 0.44% | 93.57% ± 0.45% |
| 95% nom. |  |  |  |
| $N = 60$ | 96.80% ± 0.32% | 96.50% ± 0.34% | 96.70% ± 0.33% |
| 120 | 96.90% ± 0.32% | 96.83% ± 0.32% | 97.07% ± 0.31% |
| 240 | 97.20% ± 0.30% | 96.30% ± 0.34% | 96.40% ± 0.34% |
| 480 | 97.27% ± 0.30% | 97.63% ± 0.28% | 96.77% ± 0.32% |

## References

Fidel Alfaro-Almagro, Mark Jenkinson, Neal K Bangerter, Jesper L R Andersson, Ludovica Griffanti, Gwenaëlle Douaud, Stamatios N Sotiropoulos, Saad Jbabdi, Moises Hernandez-Fernandez, Emmanuel Vallee, Diego Vidaurre, Matthew Webster, Paul McCarthy, Christopher Rorden, Alessandro Daducci, Daniel C Alexander, Hui Zhang, Iulius Dragonu, Paul M Matthews, Karla L Miller, and Stephen M Smith. Image processing and quality control for the first 10,000 brain imaging datasets from UK biobank. Neuroimage, 166:400–424, February 2018.

Deanna M Barch, Gregory C Burgess, Michael P Harms, Steven E Petersen, Bradley L Schlaggar, Maurizio Corbetta, Matthew F Glasser, Sandra Curtiss, Sachin Dixit, Cindy Feldt, Dan Nolan, Edward Bryant, Tucker Hartley, Owen Footer, James M Bjork, Russ Poldrack, Steve Smith, Heidi Johansen-Berg, Abraham Z Snyder, David C Van Essen, and WU-Minn HCP Consortium. Function in the human connectome: task-fMRI and individual differences in behavior. Neuroimage, 80:169–189, October 2013.

Gang Chen, Paul A Taylor, and Robert W Cox. Is the statistic value all we should care about in neuroimaging? Neuroimage, 147:952–959, February 2017.

Victor Chernozhukov, Denis Chetverikov, and Kengo Kato. Gaussian approximations and multiplier bootstrap for maxima of sums of high-dimensional random vectors, 2013.

Russell Davidson and Emmanuel Flachaire. The wild bootstrap, tamed at last. J. Econom., 146(1):162–169, September 2008.

Stephen A Engel and Philip C Burton. Confidence intervals for fMRI activation maps. PLoS One, 8(12):e82419, December 2013.

K J Friston, A P Holmes, K J Worsley, J P. Poline, C D Frith, and R S J Frackowiak. Statistical parametric maps in functional imaging: A general linear approach. Hum. Brain Mapp., 2(4):189–210, 1994a.

K J Friston, K J Worsley, R S Frackowiak, J C Mazziotta, and A C Evans. Assessing the significance of focal activations using their spatial extent. Hum. Brain Mapp., 1(3):210–220, 1994b.

Matthew F Glasser, Stamatios N Sotiropoulos, J Anthony Wilson, Timothy S Coalson, Bruce Fischl, Jesper L Andersson, Junqian Xu, Saad Jbabdi, Matthew Webster, Jonathan R Polimeni, David C Van Essen, Mark Jenkinson, and WU-Minn HCP Consortium. The minimal preprocessing pipelines for the human connectome project. Neuroimage, 80:105–124, October 2013.

Javier Gonzalez-Castillo, Ziad S Saad, Daniel A Handwerker, Souheil J Inati, Noah Brenowitz, and Peter A Bandettini. Whole-brain, time-locked activation with simple tasks revealed using massive averaging and model-free analysis. Proc. Natl. Acad. Sci. U. S. A., 109(14):5487–5492, April 2012.

Ahmad R Hariri, Alessandro Tessitore, Venkata S Mattay, Francesco Fera, and Daniel R Weinberger. The amygdala response to emotional stimuli: A comparison of faces and scenes. Neuroimage, 17(1):317–323, September 2002.

Paul E Meehl. Theory-Testing in psychology and physics: A methodological paradox. Philos. Sci., 34(2):103–115, June 1967.

Karla L Miller, Fidel Alfaro-Almagro, Neal K Bangerter, David L Thomas, Essa Yacoub, Junqian Xu, Andreas J Bartsch, Saad Jbabdi, Stamatios N Sotiropoulos, Jesper L R Andersson, Ludovica Griffanti, Gwenaëlle Douaud, Thomas W Okell, Peter Weale, Iulius Dragonu, Steve Garratt, Sarah Hudson, Rory Collins, Mark Jenkinson, Paul M Matthews, and Stephen M Smith. Multimodal population brain imaging in the UK biobank prospective epidemiological study. Nat. Neurosci., 19(11):1523–1536, November 2016.

Thomas E Nichols, Samir Das, Simon B Eickhoff, Alan C Evans, Tristan Glatard, Michael Hanke, Nikolaus Kriegeskorte, Michael P Milham, Russell A Poldrack, Jean-Baptiste Poline, Erika Proal, Bertrand Thirion, David C Van Essen, Tonya White, and B T Thomas Yeo. Best practices in data analysis and sharing in neuroimaging using MRI. Nat. Neurosci., 20(3):299–303, February 2017.

Regina Nuzzo. Scientific method: statistical errors. Nature, 506(7487):150–152, February 2014.

Russell A Poldrack, Jeanette A Mumford, and Thomas E Nichols. Handbook of Functional MRI Data Analysis. Cambridge University Press, August 2011.

Russell A Poldrack, Chris I Baker, Joke Durnez, Krzysztof J Gorgolewski, Paul M Matthews, Marcus R Munafò, Thomas E Nichols, Jean-Baptiste Poline, Edward Vul, and Tal Yarkoni. Scanning the horizon: towards transparent and reproducible neuroimaging research. Nat. Rev. Neurosci., 18(2):115–126, February 2017.

Jonathan D Rosenblatt, Livio Finos, Wouter D Weeda, Aldo Solari, and Jelle J Goeman. All-Resolutions inference for brain imaging. Neuroimage, 181:786–796, November 2018.

William W Rozeboom. The fallacy of the null-hypothesis significance test. Psychol. Bull., 57(5):416–428, 1960.

Stephen M Smith, Christian F Beckmann, Jesper Andersson, Edward J Auerbach, Janine Bijsterbosch, Gwenaëlle Douaud, Eugene Duff, David A Feinberg, Ludovica Griffanti, Michael P Harms, Michael Kelly, Timothy Laumann, Karla L Miller, Steen Moeller, Steve Petersen, Jonathan Power, Gholamreza Salimi-Khorshidi, Abraham Z Snyder, An T Vu, Mark W Woolrich, Junqian Xu, Essa Yacoub, Kamil Uğurbil, David C Van Essen, Matthew F Glasser, and WU-Minn HCP Consortium. Resting-state fMRI in the human connectome project. Neuroimage, 80:144–168, October 2013.

Max Sommerfeld, Stephan Sain, and Armin Schwartzman. Confidence regions for spatial excursion sets from repeated random field observations, with an application to climate. J. Am. Stat. Assoc., 113(523):1327–1340, July 2018.

F J E Telschow and A Schwartzman. Simultaneous confidence bands for functional data using the gaussian kinematic formula. arXiv preprint arXiv:1901.06386, 2019.

K UIudag, B Müller-Bierl, and K Ugurbil. An integrative model for neuronal activity-induced signal changes for gradient and spin echo functional imaging, 2009.

Ronald L Wasserstein, Nicole A Lazar, and Others. The ASA’s statement on p-values: context, process, and purpose. Am. Stat., 70(2):129–133, 2016.

Anderson M Winkler, Gerard R Ridgway, Gwenaëlle Douaud, Thomas E Nichols, and Stephen M Smith. Faster permutation inference in brain imaging. Neuroimage, 141:502–516, November 2016.

Choong-Wan Woo, Anjali Krishnan, and Tor D Wager. Cluster-extent based thresholding in fMRI analyses: pitfalls and recommendations. Neuroimage, 91:412–419, May 2014.

